## Supplementary figures and images for "Human cytomegalovirus infection coopts chromatin organization to diminish TEAD1 transcription factor activity"

### Supplemental Figure 1

**Supplemental Figure S1**

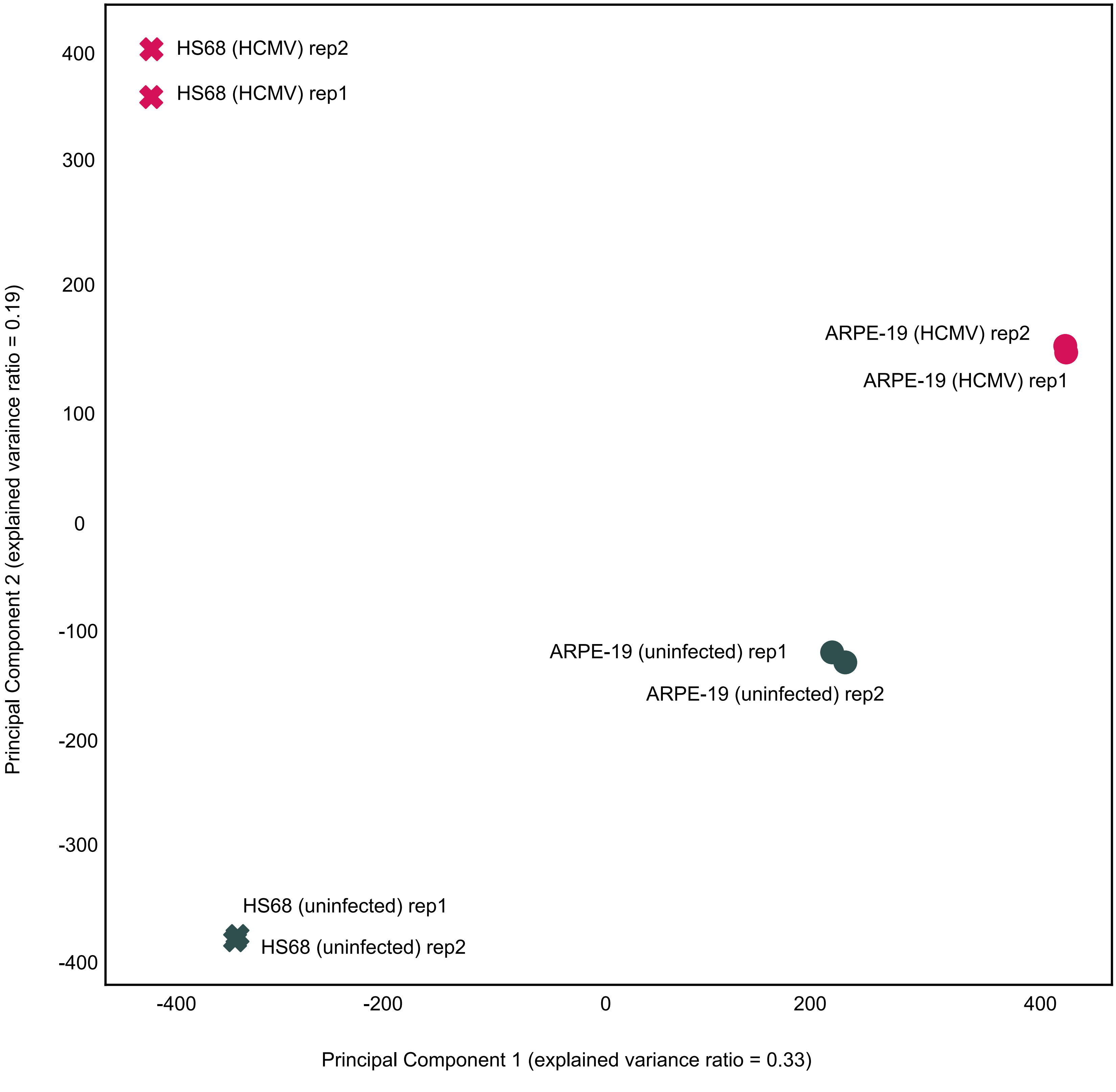

### Supplemental Figure 2

##### Supplemental Figure S2

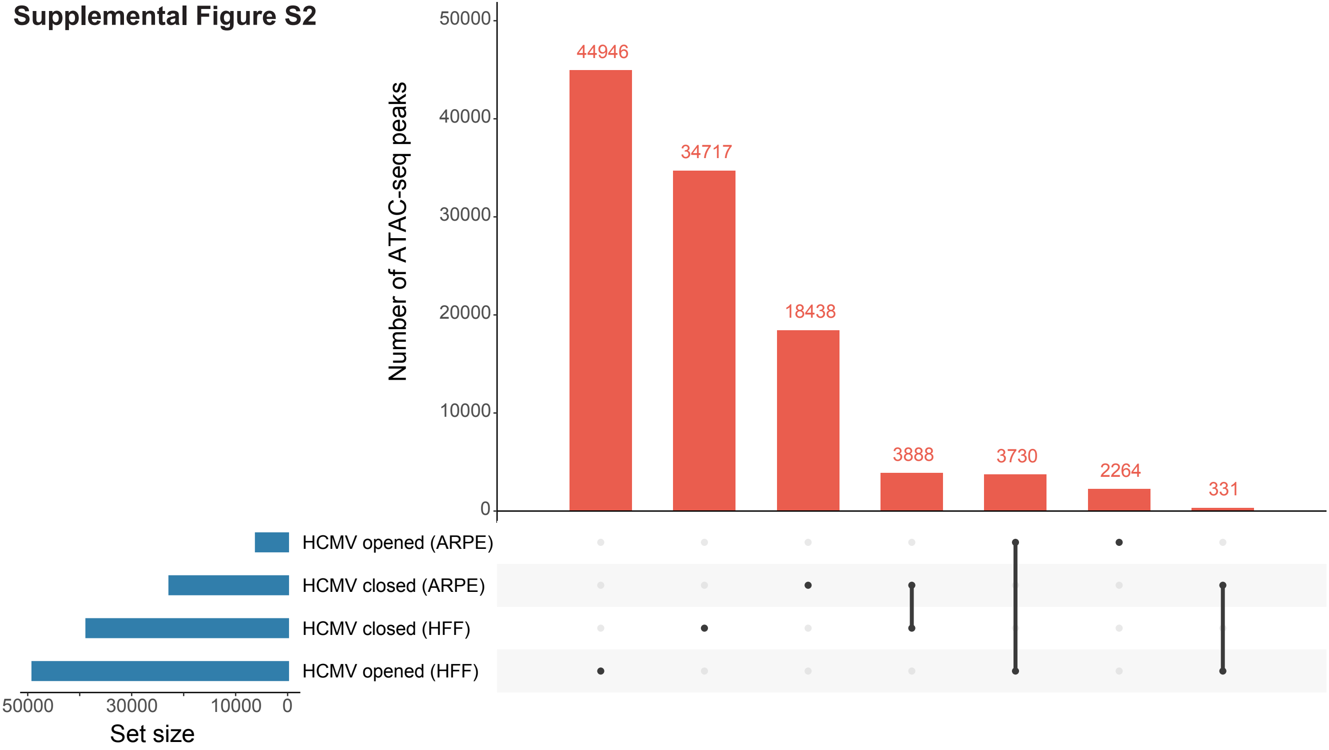

### Supplemental Figure 4

**Supplemental Figure S4**

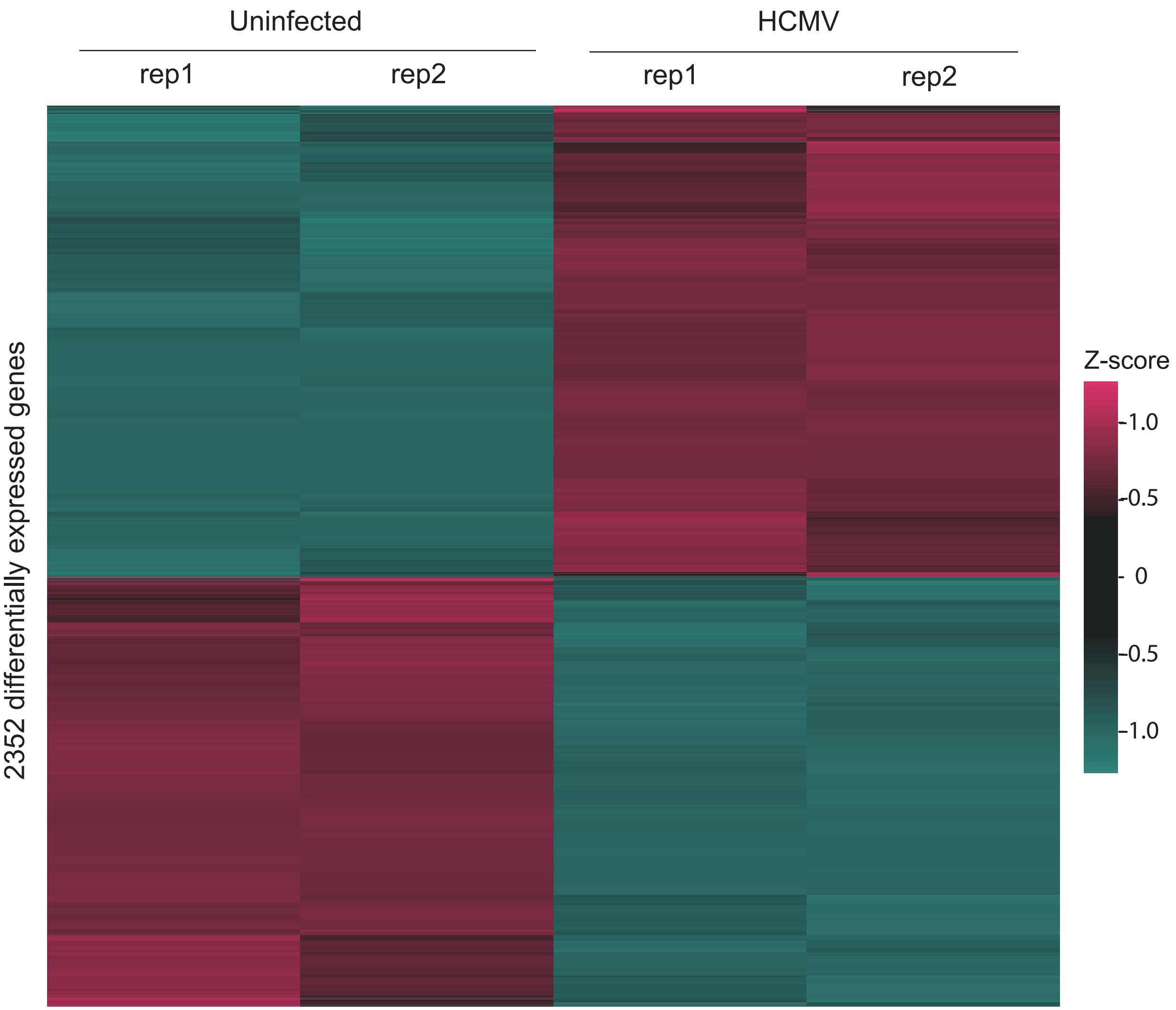

### Supplemental Figure 5

**Supplemental Figure S5** : Western blots of uninfected and HCMV infected fibroblasts

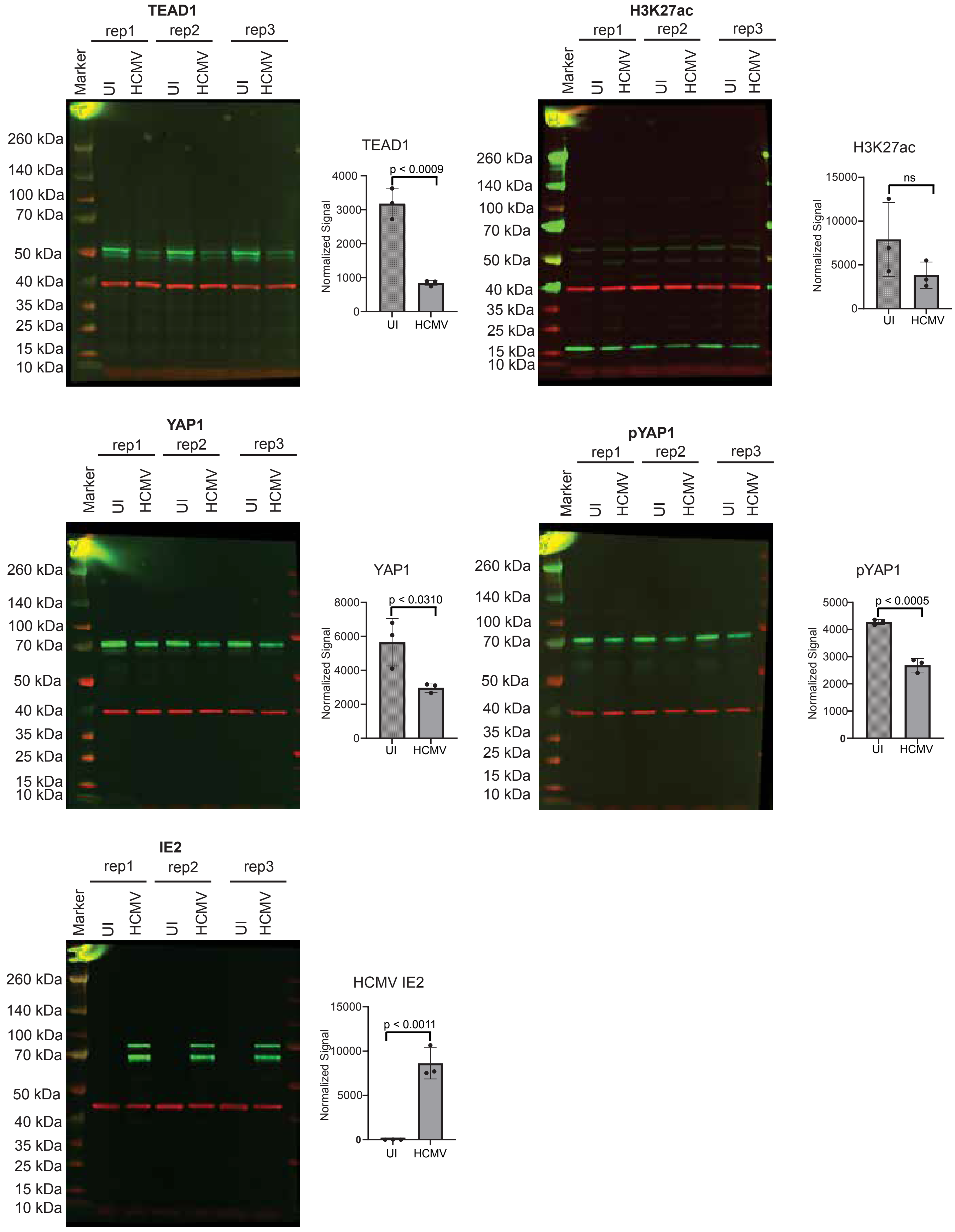

### Supplemental Figure 7

#### Supplemental Figure S7

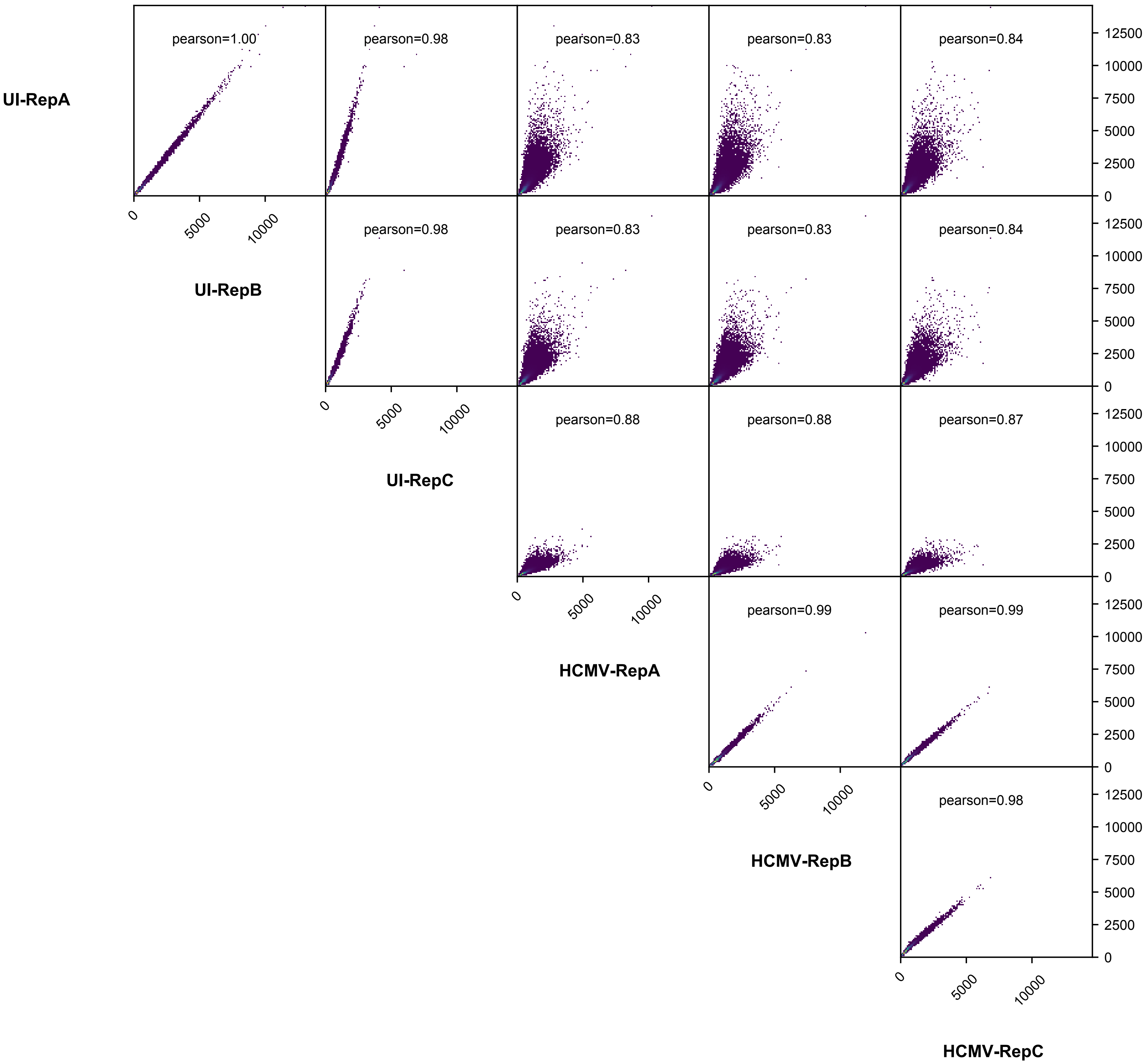

### Supplemental Figure 8

**Supplemental Figure S8**

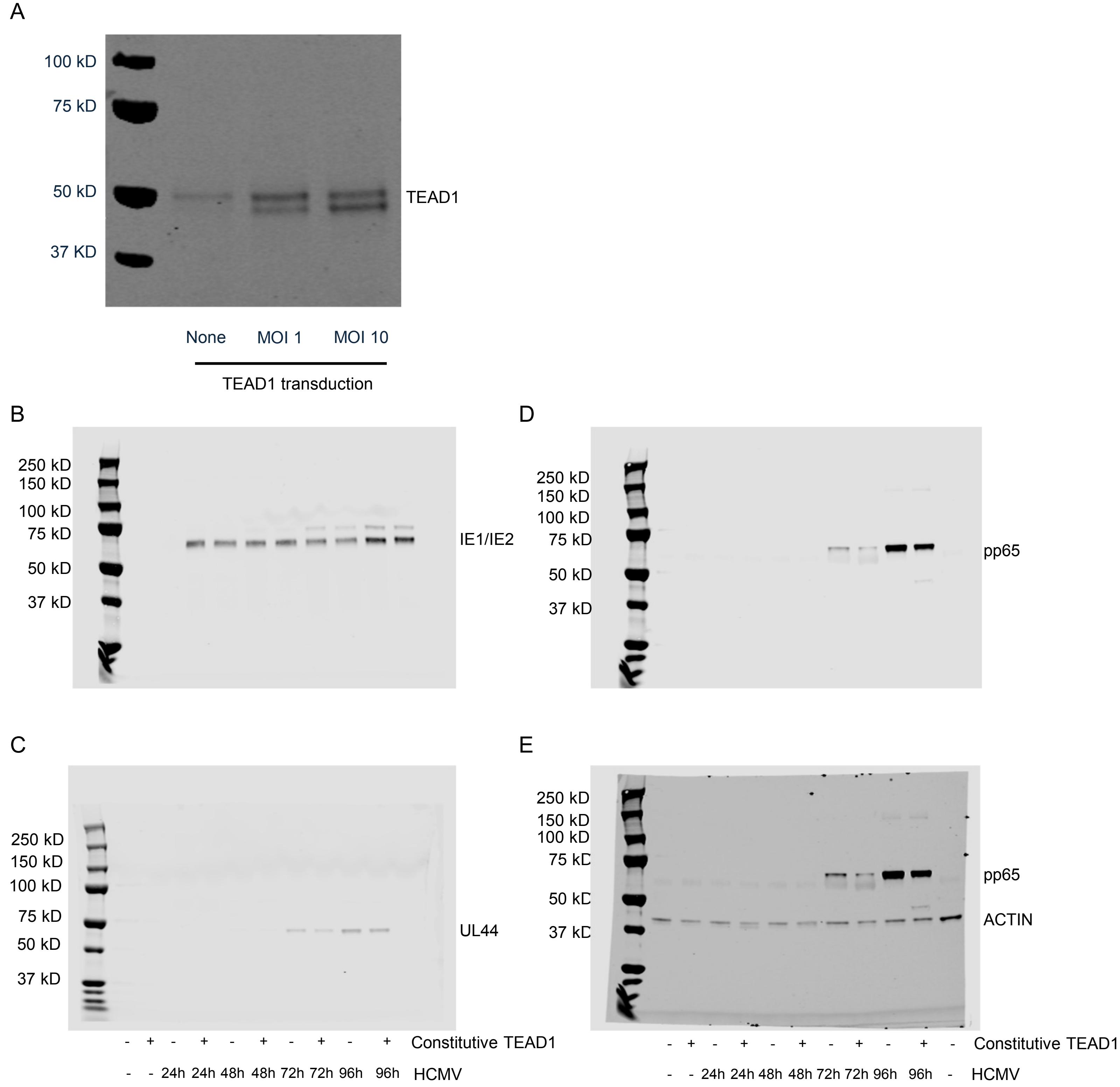
