## Supplemental Figure 3 for "Human cytomegalovirus infection coopts chromatin organization to diminish TEAD1 transcription factor activity"

**Supplemental Figure S3 :** Heatmap of ChIP-seq peaks for TEAD1, CTCF, and H3K27ac between replicates in uninfected and HCMV infected fibroblasts

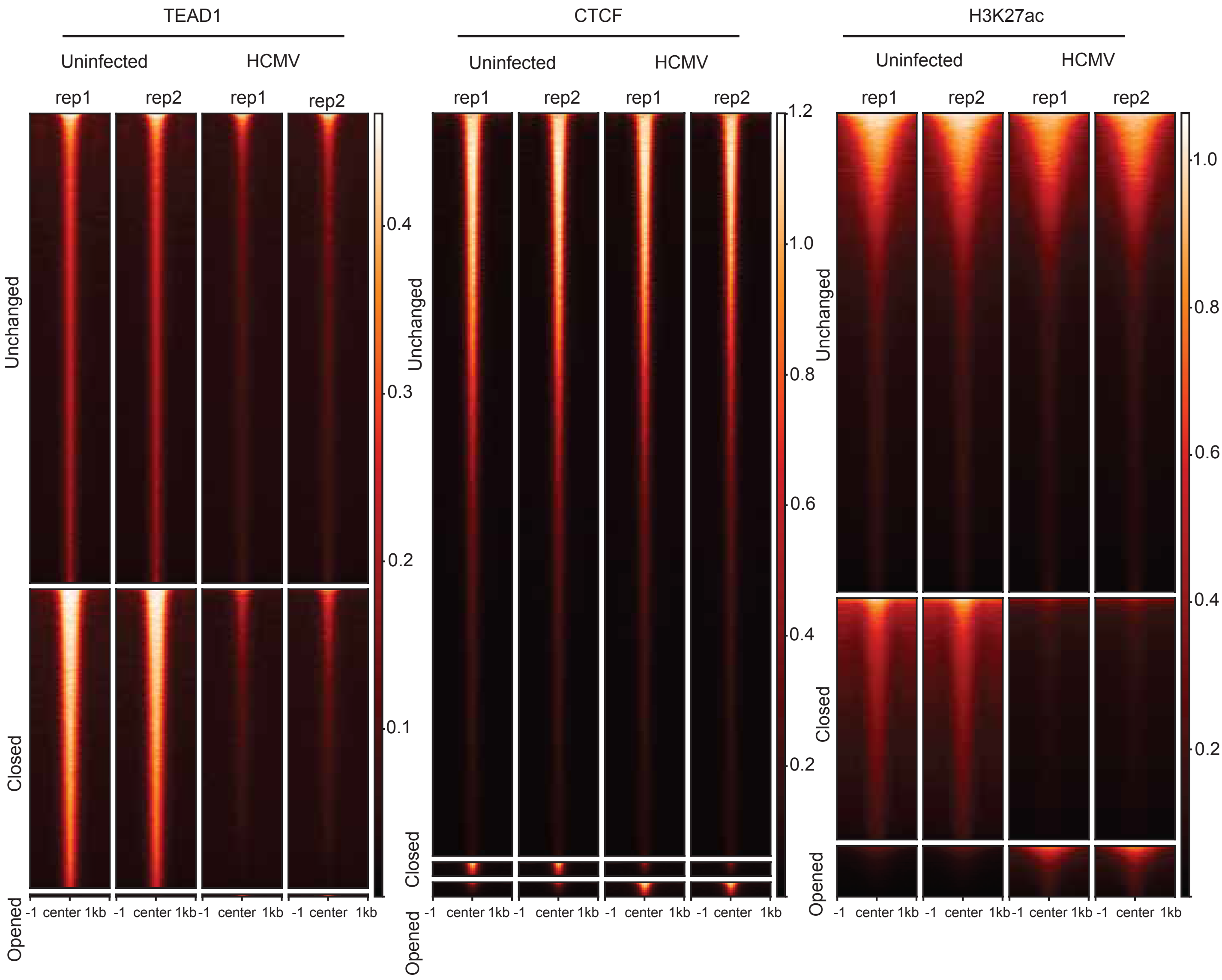
