## Supplemental Figure 6 for "Human cytomegalovirus infection coopts chromatin organization to diminish TEAD1 transcription factor activity"

**Supplemental Figure S6**

A. Splicing counts

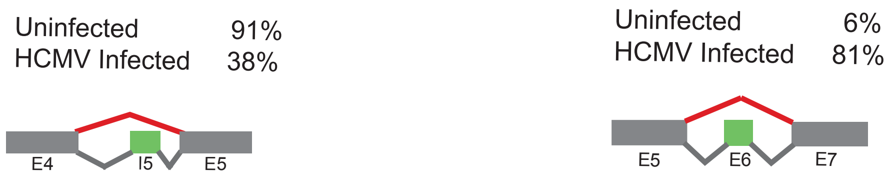


B. Sashimi plots

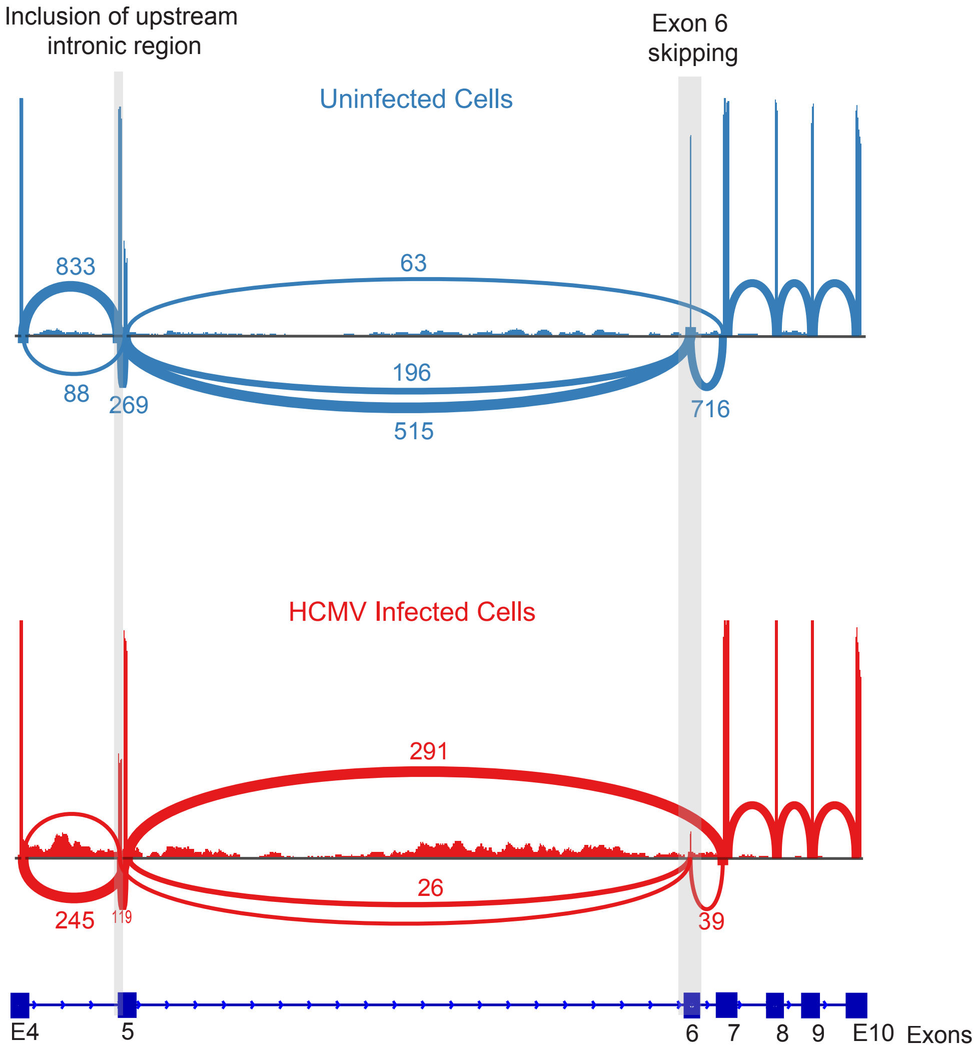


C. TEAD1 sequence showing PCR primers and exon 6 sequence

```
AAATCTCGTGATTTTCATTCCAAGCTAAAGGTAACAAGCATGGATCAGACTGCAAAGGATAAGGCCCTGCAGCACATGGCGGCCATGTCCTCAGCCCAGA
---------+---------+---------+---------+---------+---------+---------+---------+---------+---------+
TTTAGAGCACTAAAAGTAAGGTTCGATTTCCATTGTTCGTACCTAGTCTGACGTTTCCTATTCCGGGACGTCGTGTACCGCCGGTACAGGAGTCGGGTCT
K  S  R  D  F  H  S  K  L  K  V  T  S  M  D  Q  T  A  K  D  K  A  L  Q  H  M  A  A  M  S  S  A  Q  I
```
