## Supplementary material for "Human cytomegalovirus infection coopts chromatin organization to diminish TEAD1 transcription factor activity": html files for QC: HFF_deseq.html

Preliminary DESeq2 Results


### Preliminary DESeq2 Results

###### 02 December, 2022

- Run Summary
  - Analysis Parameters
- DESeq2 Processing Information
  - PCA
  - Dispersion Estimate
  - Sparsity
- DEG Comparisons
  - cmv\_status: TB40E vs UI
  - cmv\_status: UI vs TB40E
- Explanation of files
- Environment Information
  - Parameters used
  - Packages
  - Platform Info
  - Code

**NOTE**: This report is considered to be a first pass at differential expression. There are a lot of variables to consider and many moving parts to this document, so please consider verifying anything you find interesting independently.

#### Run Summary

| Sample Information | | | |
| --- | --- | --- | --- |
|  | cell\_line | cmv\_status | quant\_sf |
| --- | --- | --- | --- |
| HS68\_UI\_rep1 | HS68 | UI | /data/weirauchlab/project/CMV/datasets/rnaseq/tb40e/nf-core\_ver2/hcmv/analysis/star\_salmon/HS68\_UI\_rep1/quant.sf |
| HS68\_UI\_rep2 | HS68 | UI | /data/weirauchlab/project/CMV/datasets/rnaseq/tb40e/nf-core\_ver2/hcmv/analysis/star\_salmon/HS68\_UI\_rep2/quant.sf |
| HS68\_HCMV\_rep1 | HS68 | TB40E | /data/weirauchlab/project/CMV/datasets/rnaseq/tb40e/nf-core\_ver2/hcmv/analysis/star\_salmon/HS68\_TB40E-GFP\_rep1/quant.sf |
| HS68\_HCMV\_rep2 | HS68 | TB40E | /data/weirauchlab/project/CMV/datasets/rnaseq/tb40e/nf-core\_ver2/hcmv/analysis/star\_salmon/HS68\_TB40E-GFP\_rep2/quant.sf |

##### Analysis Parameters

| Summary of DESeq2 Inputs | |
| --- | --- |
| DESeqDataSet Preparation | |
| Design File | sample\_summary.csv |
| Counts Format | salmon |
| Column with counts files | quant\_sf |
| Comparison Conditions | |
| Design Formula | ~cmv\_status |
| Contrast Column | cmv\_status |

#### DESeq2 Processing Information

##### PCA

##### Dispersion Estimate

```
## png 
##   2
```

```
## png 
##   2
```

##### Sparsity

```
## png 
##   2
```

```
## png 
##   2
```

#### DEG Comparisons

| Schema for classifying gene status | | | | | |
| --- | --- | --- | --- | --- | --- |
|  | baseMean | log2FoldChange | pvalue | padj | Notes |
| --- | --- | --- | --- | --- | --- |
| Zero Count | = 0 | NA | NA | NA | Zero count genes are removed from consideration prior to statistical analysis. |
| Outlier | > 0 | Exists | NA | NA | The gene is considered to be an outlier and is censured from analysis. |
| Low Count | > 0 | Exists | Exists | NA | An independent filtering step is performed prior to analysis that tries to establish a threshold for 'low count' genes. DESeq2's IF step tries to maximize the number of DEGs, so this may vary between contrasts. |
| Insignificant | > 0 | Exists | Exists | >= 0.01 | These are part of the actual analysis |
| DEG (Differentially Expressed Gene) | > 0 | \|log2FC\| > 1 | Exists | < 0.01 | These are the genes that are considered to be of interest. |

**Use the tabs to select a comparison.**

##### cmv\_status: TB40E vs UI

Summary of the classification for each gene after performing DESeq2 analysis.

| DESeq2 Results Summary | | |
| --- | --- | --- |
| cmv\_status: TB40E vs UI | | |
|  | Number | Percentage |
| --- | --- | --- |
| DEG | 2,511 | 4.33% |
| Insignificant | 15,580 | 26.87% |
| Low Count | 8,070 | 13.92% |
| Zero Count | 31,826 | 54.88% |
| Total | 57,987 | 100.00% |

Top 10 up/down-regulated genes relative to control.

| Top Differentially Expressed Genes | | | | | | | | |
| --- | --- | --- | --- | --- | --- | --- | --- | --- |
| cmv\_status: TB40E vs UI | | | | | | | | |
|  | gene\_name | gene\_type | baseMean | log2FoldChange | lfcSE | stat | pvalue | padj |
| --- | --- | --- | --- | --- | --- | --- | --- | --- |
| Upregulated | | | | | | | | |
| ENSG00000025039.10 | RRAGD | protein\_coding | 3,361.354 | 8.338 | 0.219 | 33.481 | 9.04 × 10^−246^ | 3.80 × 10^−243^ |
| ENSG00000169429.6 | IL8 | protein\_coding | 5,204.774 | 5.275 | 0.144 | 29.787 | 5.74 × 10^−195^ | 1.96 × 10^−192^ |
| UL40 | UL40 | CMV | 100,191.024 | 15.826 | 0.519 | 28.547 | 3.05 × 10^−179^ | 9.34 × 10^−177^ |
| ENSG00000166145.10 | SPINT1 | protein\_coding | 2,135.812 | 7.123 | 0.216 | 28.317 | 2.16 × 10^−176^ | 6.39 × 10^−174^ |
| US28 | US28 | CMV | 92,056.276 | 15.712 | 0.521 | 28.235 | 2.19 × 10^−175^ | 6.38 × 10^−173^ |
| UL5 | UL5 | CMV | 81,164.729 | 15.546 | 0.523 | 27.792 | 5.35 × 10^−170^ | 1.45 × 10^−167^ |
| US18 | US18 | CMV | 75,623.345 | 15.454 | 0.525 | 27.555 | 3.84 × 10^−167^ | 1.02 × 10^−164^ |
| UL31 | UL31 | CMV | 67,926.080 | 15.312 | 0.527 | 27.183 | 1.02 × 10^−162^ | 2.54 × 10^−160^ |
| UL44 | UL44 | CMV | 66,024.604 | 15.275 | 0.527 | 27.084 | 1.53 × 10^−161^ | 3.70 × 10^−159^ |
| UL49 | UL49 | CMV | 65,055.245 | 15.256 | 0.527 | 27.035 | 5.67 × 10^−161^ | 1.35 × 10^−158^ |
| Downregulated | | | | | | | | |
| ENSG00000114251.9 | WNT5A | protein\_coding | 62,005.021 | −3.976 | 0.069 | −43.145 | 0.00 | 0.00 |
| ENSG00000105974.7 | CAV1 | protein\_coding | 32,851.779 | −3.986 | 0.079 | −37.821 | 0.00 | 0.00 |
| ENSG00000011465.12 | DCN | protein\_coding | 65,930.818 | −4.003 | 0.074 | −40.339 | 0.00 | 0.00 |
| ENSG00000075223.9 | SEMA3C | protein\_coding | 23,738.750 | −4.219 | 0.079 | −40.754 | 0.00 | 0.00 |
| ENSG00000113083.8 | LOX | protein\_coding | 28,351.738 | −4.275 | 0.070 | −46.851 | 0.00 | 0.00 |
| ENSG00000112902.7 | SEMA5A | protein\_coding | 12,260.734 | −4.327 | 0.086 | −38.606 | 0.00 | 0.00 |
| ENSG00000115380.14 | EFEMP1 | protein\_coding | 16,610.603 | −4.728 | 0.085 | −44.101 | 0.00 | 0.00 |
| ENSG00000065534.14 | MYLK | protein\_coding | 26,233.889 | −4.895 | 0.076 | −51.571 | 0.00 | 0.00 |
| ENSG00000108821.9 | COL1A1 | protein\_coding | 23,251.751 | −5.104 | 0.105 | −39.016 | 0.00 | 0.00 |
| ENSG00000107796.8 | ACTA2 | protein\_coding | 3,760.599 | −6.018 | 0.133 | −37.754 | 0.00 | 0.00 |
|  |  |  |  |  |  |  |  |  |
| --- | --- | --- | --- | --- | --- | --- | --- | --- |
| **baseMean**: The mean of the normalized counts between all samples | | | | | | | | |
| **log2FoldChange**: log_2_ fold change of the target / control | | | | | | | | |
| **lfcSE**: Standard error of the log_2_ fold change | | | | | | | | |
| **stat**: The statistic used to calculate the p-value | | | | | | | | |
| **pvalue**: p-value of the comparison, prior to adjustment | | | | | | | | |
| **padj**: The adjusted p-value, accounting for multiple comparisons | | | | | | | | |

###### Volcano Plot

###### MA Plot

###### Independent Filtering Plot

###### Full DEG Heatmap (all samples)

###### Filtered DEG Heatmap (only contrast samples)

##### cmv\_status: UI vs TB40E

Summary of the classification for each gene after performing DESeq2 analysis.

| DESeq2 Results Summary | | |
| --- | --- | --- |
| cmv\_status: UI vs TB40E | | |
|  | Number | Percentage |
| --- | --- | --- |
| DEG | 2,511 | 4.33% |
| Insignificant | 15,580 | 26.87% |
| Low Count | 8,070 | 13.92% |
| Zero Count | 31,826 | 54.88% |
| Total | 57,987 | 100.00% |

Top 10 up/down-regulated genes relative to control.

| Top Differentially Expressed Genes | | | | | | | | |
| --- | --- | --- | --- | --- | --- | --- | --- | --- |
| cmv\_status: UI vs TB40E | | | | | | | | |
|  | gene\_name | gene\_type | baseMean | log2FoldChange | lfcSE | stat | pvalue | padj |
| --- | --- | --- | --- | --- | --- | --- | --- | --- |
| Upregulated | | | | | | | | |
| ENSG00000107796.8 | ACTA2 | protein\_coding | 3,760.599 | 6.018 | 0.133 | 37.754 | 0.00 | 0.00 |
| ENSG00000108821.9 | COL1A1 | protein\_coding | 23,251.751 | 5.104 | 0.105 | 39.016 | 0.00 | 0.00 |
| ENSG00000065534.14 | MYLK | protein\_coding | 26,233.889 | 4.895 | 0.076 | 51.571 | 0.00 | 0.00 |
| ENSG00000115380.14 | EFEMP1 | protein\_coding | 16,610.603 | 4.728 | 0.085 | 44.101 | 0.00 | 0.00 |
| ENSG00000112902.7 | SEMA5A | protein\_coding | 12,260.734 | 4.327 | 0.086 | 38.606 | 0.00 | 0.00 |
| ENSG00000113083.8 | LOX | protein\_coding | 28,351.738 | 4.275 | 0.070 | 46.851 | 0.00 | 0.00 |
| ENSG00000075223.9 | SEMA3C | protein\_coding | 23,738.750 | 4.219 | 0.079 | 40.754 | 0.00 | 0.00 |
| ENSG00000011465.12 | DCN | protein\_coding | 65,930.818 | 4.003 | 0.074 | 40.339 | 0.00 | 0.00 |
| ENSG00000105974.7 | CAV1 | protein\_coding | 32,851.779 | 3.986 | 0.079 | 37.821 | 0.00 | 0.00 |
| ENSG00000114251.9 | WNT5A | protein\_coding | 62,005.021 | 3.976 | 0.069 | 43.145 | 0.00 | 0.00 |
| Downregulated | | | | | | | | |
| ENSG00000025039.10 | RRAGD | protein\_coding | 3,361.354 | −8.338 | 0.219 | −33.481 | 9.04 × 10^−246^ | 3.80 × 10^−243^ |
| ENSG00000169429.6 | IL8 | protein\_coding | 5,204.774 | −5.275 | 0.144 | −29.787 | 5.74 × 10^−195^ | 1.96 × 10^−192^ |
| UL40 | UL40 | CMV | 100,191.024 | −15.826 | 0.519 | −28.547 | 3.05 × 10^−179^ | 9.34 × 10^−177^ |
| ENSG00000166145.10 | SPINT1 | protein\_coding | 2,135.812 | −7.123 | 0.216 | −28.317 | 2.16 × 10^−176^ | 6.39 × 10^−174^ |
| US28 | US28 | CMV | 92,056.276 | −15.712 | 0.521 | −28.235 | 2.19 × 10^−175^ | 6.38 × 10^−173^ |
| UL5 | UL5 | CMV | 81,164.729 | −15.546 | 0.523 | −27.792 | 5.35 × 10^−170^ | 1.45 × 10^−167^ |
| US18 | US18 | CMV | 75,623.345 | −15.454 | 0.525 | −27.555 | 3.84 × 10^−167^ | 1.02 × 10^−164^ |
| UL31 | UL31 | CMV | 67,926.080 | −15.312 | 0.527 | −27.183 | 1.02 × 10^−162^ | 2.54 × 10^−160^ |
| UL44 | UL44 | CMV | 66,024.604 | −15.275 | 0.527 | −27.084 | 1.53 × 10^−161^ | 3.70 × 10^−159^ |
| UL49 | UL49 | CMV | 65,055.245 | −15.256 | 0.527 | −27.035 | 5.67 × 10^−161^ | 1.35 × 10^−158^ |
|  |  |  |  |  |  |  |  |  |
| --- | --- | --- | --- | --- | --- | --- | --- | --- |
| **baseMean**: The mean of the normalized counts between all samples | | | | | | | | |
| **log2FoldChange**: log_2_ fold change of the target / control | | | | | | | | |
| **lfcSE**: Standard error of the log_2_ fold change | | | | | | | | |
| **stat**: The statistic used to calculate the p-value | | | | | | | | |
| **pvalue**: p-value of the comparison, prior to adjustment | | | | | | | | |
| **padj**: The adjusted p-value, accounting for multiple comparisons | | | | | | | | |

###### Volcano Plot

###### MA Plot

###### Independent Filtering Plot

###### Full DEG Heatmap (all samples)

###### Filtered DEG Heatmap (only contrast samples)

#### Explanation of files

There are several files that are exported with an analysis.

Within your main results dir, **deseq2\_results\_stringent**, you will have a few files, as well as folders for any contrast that was specified.

| File Type | File Pattern | Explanation |
| --- | --- | --- |
| DESeq2 data object | DESeq2\_object.RDS | An R-loadable object containing the object created by the DESeq() function. |
| DESeq2 VST counts | vst\_counts.csv | The variance-stabilized transformation counts output after using the DESeq() and vst() functions. |
| DESeq2 normalized counts | normalized\_counts.csv | The normalized counts output after using the DESeq() and counts(normalized = TRUE) functions. |
| Dispersion plot | DESeq2\_dispersion\_plot.(png\|pdf) | The dispersion plot output by DESeq2. |
| Sparsity plot | DESeq2\_sparsity\_plot.(png\|pdf) | The sparsity plot output by DESeq2. |
| PCA plot | DESeq2\_pca\_plot.(png\|pdf) | The PCA plot output by DESeq2. |
| DEG Heatmap | <contrast folder>/<contrast>\_deg\_heatmap\_plot.(png\|pdf) | A heatmap of z-scored vst counts for the differentially expressed genes in the contrast. |
| Independent Filtering plot | <contrast folder>/<contrast>\_independent-filter\_plot.(png\|pdf) | The independent filtering plot for the contrast. |
| MA plot | <contrast folder>/<contrast>\_ma\_plot.(png\|pdf) | The MA plot for gene expression values in the contrast. |
| Volcano plot | <contrast folder>/<contrast>\_ma\_plot.(png\|pdf) | The volcano plot for gene expression values in the contrast. |
| DESeq2 results object | <contrast folder>/<contrast>\_results.rds | The results object calculated by DESeq2 for the contrast. |
| DESeq2 results csv | <contrast folder>/<contrast>\_results.csv | A csv file containing the results calculated by DESeq2 for the contrast. The classification column annotates gene status according to DESeq2's methodology. |
| DESeq2 results xlsx | <contrast folder>/<contrast>\_results.xlsx | An xlsx file containing the results calculated by DESeq2 for the contrast. Each tab contains the genes matching the classification according to DESeq2's methodology. |

#### Environment Information

##### Parameters used

| parameter | value |
| --- | --- |
| counts\_file |  |
| contrasts |  |
| colData\_scrub\_filters | short\_name, read1, read2, experimental\_id, id, quant\_sf |
| annotation\_file |  |
| tximport\_params |  |
| DESeqDataSetFromTximport\_params |  |
| DESeq\_params | betaPrior = TRUE |
| DEG\_heatmap\_min\_color | #00FFFF |
| DEG\_heatmap\_mid\_color | #000000 |
| DEG\_heatmap\_max\_color | #E10022 |
| DEG\_heatmap\_cluster\_rows | TRUE |
| DEG\_heatmap\_cluster\_cols | TRUE |
| DEG\_heatmap\_show\_row\_names | FALSE |
| DEG\_heatmap\_show\_col\_names | FALSE |
| pca\_groups |  |
| volcano\_plot\_colors | DEG = #E10022, Insignificant = #000000, Low Count = #00FF00, Zero Count = #0000FF |
| ma\_plot\_colors | DEG = #E10022, Insignificant = #000000, Low Count = #00FF00, Zero Count = #0000FF |
| volcano\_plot\_export | width = 7, height = 7, units = in, dpi = 300 |
| ma\_plot\_export | width = 7, height = 7, units = in, dpi = 300 |
| if\_plot\_export | width = 7, height = 7, units = in, dpi = 300 |
| heatmap\_export | width = 7, height = 7, units = in, res = 300 |
| pca\_plot\_export | width = 7, height = 7, units = in, dpi = 300 |
| dispersion\_export | width = 7, height = 7, units = in, res = 300 |
| sparsity\_export | width = 7, height = 7, units = in, res = 300 |
| version | 1.1 |
| run\_volcano | TRUE |
| run\_ma | TRUE |
| run\_if | TRUE |
| run\_heatmap | TRUE |
| run\_filtered\_heatmap | TRUE |
| counts\_rowname | Geneid |
| design\_file | sample\_summary.csv |
| sample\_id\_col | id |
| design\_formula | ~cmv\_status |
| contrast\_col | cmv\_status |
| gtf\_file | /data/weirauchlab/databank/genome/nfcore/cmv/hg19\_TB40-BAC4.gtf |
| annotation\_merge\_col | gene\_id |
| results\_dir | deseq2\_results\_stringent |
| counts\_format | salmon |
| sf\_file\_col | quant\_sf |
| DESeq\_results\_params | alpha = 0.01, lfcThreshold = 1, altHypothesis = greaterAbs |

##### Packages

| package | ondiskversion | loadedversion | path | loadedpath | attached | is\_base | date | source | md5ok | library |
| --- | --- | --- | --- | --- | --- | --- | --- | --- | --- | --- |
| Biobase | 2.50.0 | 2.50.0 | /usr/local/R/4.0.2/lib64/R/library/Biobase | /usr/local/R/4.0.2/lib64/R/library/Biobase | TRUE | FALSE | 2020-10-27 | Bioconductor | NA | /usr/local/R/4.0.2/lib64/R/library |
| BiocGenerics | 0.38.0 | 0.38.0 | /usr/local/R/4.0.2/lib64/R/library/BiocGenerics | /usr/local/R/4.0.2/lib64/R/library/BiocGenerics | TRUE | FALSE | 2021-05-19 | Bioconductor | NA | /usr/local/R/4.0.2/lib64/R/library |
| circlize | 0.4.12 | 0.4.12 | /usr/local/R/4.0.2/lib64/R/library/circlize | /usr/local/R/4.0.2/lib64/R/library/circlize | TRUE | FALSE | 2021-01-08 | CRAN (R 4.0.2) | NA | /usr/local/R/4.0.2/lib64/R/library |
| ComplexHeatmap | 2.6.2 | 2.6.2 | /usr/local/R/4.0.2/lib64/R/library/ComplexHeatmap | /usr/local/R/4.0.2/lib64/R/library/ComplexHeatmap | TRUE | FALSE | 2020-11-12 | Bioconductor | NA | /usr/local/R/4.0.2/lib64/R/library |
| DESeq2 | 1.30.1 | 1.30.1 | /usr/local/R/4.0.2/lib64/R/library/DESeq2 | /usr/local/R/4.0.2/lib64/R/library/DESeq2 | TRUE | FALSE | 2021-02-19 | Bioconductor | NA | /usr/local/R/4.0.2/lib64/R/library |
| dplyr | 1.0.8 | 1.0.8 | /usr/local/R/4.0.2/lib64/R/library/dplyr | /usr/local/R/4.0.2/lib64/R/library/dplyr | TRUE | FALSE | 2022-02-08 | CRAN (R 4.0.2) | NA | /usr/local/R/4.0.2/lib64/R/library |
| forcats | 0.5.1 | 0.5.1 | /usr/local/R/4.0.2/lib64/R/library/forcats | /usr/local/R/4.0.2/lib64/R/library/forcats | TRUE | FALSE | 2021-01-27 | CRAN (R 4.0.2) | NA | /usr/local/R/4.0.2/lib64/R/library |
| GenomeInfoDb | 1.28.4 | 1.28.4 | /usr/local/R/4.0.2/lib64/R/library/GenomeInfoDb | /usr/local/R/4.0.2/lib64/R/library/GenomeInfoDb | TRUE | FALSE | 2021-09-05 | Bioconductor | NA | /usr/local/R/4.0.2/lib64/R/library |
| GenomicRanges | 1.42.0 | 1.42.0 | /usr/local/R/4.0.2/lib64/R/library/GenomicRanges | /usr/local/R/4.0.2/lib64/R/library/GenomicRanges | TRUE | FALSE | 2020-10-27 | Bioconductor | NA | /usr/local/R/4.0.2/lib64/R/library |
| ggplot2 | 3.3.5 | 3.3.5 | /usr/local/R/4.0.2/lib64/R/library/ggplot2 | /usr/local/R/4.0.2/lib64/R/library/ggplot2 | TRUE | FALSE | 2021-06-25 | CRAN (R 4.0.2) | NA | /usr/local/R/4.0.2/lib64/R/library |
| gt | 0.5.0 | 0.5.0 | /users/parz1z/R/x86\_64-pc-linux-gnu-library/4.0/gt | /users/parz1z/R/x86\_64-pc-linux-gnu-library/4.0/gt | TRUE | FALSE | 2022-04-21 | CRAN (R 4.0.2) | NA | /users/parz1z/R/x86\_64-pc-linux-gnu-library/4.0 |
| IRanges | 2.24.1 | 2.24.1 | /usr/local/R/4.0.2/lib64/R/library/IRanges | /usr/local/R/4.0.2/lib64/R/library/IRanges | TRUE | FALSE | 2020-12-12 | Bioconductor | NA | /usr/local/R/4.0.2/lib64/R/library |
| knitr | 1.37 | 1.37 | /usr/local/R/4.0.2/lib64/R/library/knitr | /usr/local/R/4.0.2/lib64/R/library/knitr | TRUE | FALSE | 2021-12-16 | CRAN (R 4.0.2) | NA | /usr/local/R/4.0.2/lib64/R/library |
| MatrixGenerics | 1.7.0 | 1.7.0 | /usr/local/R/4.0.2/lib64/R/library/MatrixGenerics | /usr/local/R/4.0.2/lib64/R/library/MatrixGenerics | TRUE | FALSE | 2022-03-03 | Github (Bioconductor/MatrixGenerics@dfc3746) | NA | /usr/local/R/4.0.2/lib64/R/library |
| matrixStats | 0.61.0 | 0.61.0 | /usr/local/R/4.0.2/lib64/R/library/matrixStats | /usr/local/R/4.0.2/lib64/R/library/matrixStats | TRUE | FALSE | 2021-09-17 | CRAN (R 4.0.2) | NA | /usr/local/R/4.0.2/lib64/R/library |
| openxlsx | 4.2.3 | 4.2.3 | /usr/local/R/4.0.2/lib64/R/library/openxlsx | /usr/local/R/4.0.2/lib64/R/library/openxlsx | TRUE | FALSE | 2020-10-27 | CRAN (R 4.0.2) | NA | /usr/local/R/4.0.2/lib64/R/library |
| purrr | 0.3.4 | 0.3.4 | /usr/local/R/4.0.2/lib64/R/library/purrr | /usr/local/R/4.0.2/lib64/R/library/purrr | TRUE | FALSE | 2020-04-17 | CRAN (R 4.0.2) | NA | /usr/local/R/4.0.2/lib64/R/library |
| readr | 1.4.0 | 1.4.0 | /usr/local/R/4.0.2/lib64/R/library/readr | /usr/local/R/4.0.2/lib64/R/library/readr | TRUE | FALSE | 2020-10-05 | CRAN (R 4.0.2) | NA | /usr/local/R/4.0.2/lib64/R/library |
| S4Vectors | 0.28.1 | 0.28.1 | /usr/local/R/4.0.2/lib64/R/library/S4Vectors | /usr/local/R/4.0.2/lib64/R/library/S4Vectors | TRUE | FALSE | 2020-12-09 | Bioconductor | NA | /usr/local/R/4.0.2/lib64/R/library |
| stringr | 1.4.0 | 1.4.0 | /usr/local/R/4.0.2/lib64/R/library/stringr | /usr/local/R/4.0.2/lib64/R/library/stringr | TRUE | FALSE | 2019-02-10 | CRAN (R 4.0.2) | NA | /usr/local/R/4.0.2/lib64/R/library |
| SummarizedExperiment | 1.20.0 | 1.20.0 | /usr/local/R/4.0.2/lib64/R/library/SummarizedExperiment | /usr/local/R/4.0.2/lib64/R/library/SummarizedExperiment | TRUE | FALSE | 2020-10-27 | Bioconductor | NA | /usr/local/R/4.0.2/lib64/R/library |
| tibble | 3.1.6 | 3.1.6 | /usr/local/R/4.0.2/lib64/R/library/tibble | /usr/local/R/4.0.2/lib64/R/library/tibble | TRUE | FALSE | 2021-11-07 | CRAN (R 4.0.2) | NA | /usr/local/R/4.0.2/lib64/R/library |
| tidyr | 1.2.0 | 1.2.0 | /usr/local/R/4.0.2/lib64/R/library/tidyr | /usr/local/R/4.0.2/lib64/R/library/tidyr | TRUE | FALSE | 2022-02-01 | CRAN (R 4.0.2) | NA | /usr/local/R/4.0.2/lib64/R/library |
| tidyverse | 1.3.1 | 1.3.1 | /usr/local/R/4.0.2/lib64/R/library/tidyverse | /usr/local/R/4.0.2/lib64/R/library/tidyverse | TRUE | FALSE | 2021-04-15 | CRAN (R 4.0.2) | NA | /usr/local/R/4.0.2/lib64/R/library |
| tximport | 1.18.0 | 1.18.0 | /usr/local/R/4.0.2/lib64/R/library/tximport | /usr/local/R/4.0.2/lib64/R/library/tximport | TRUE | FALSE | 2020-10-27 | Bioconductor | NA | /usr/local/R/4.0.2/lib64/R/library |

##### Platform Info

| Metric | Value |
| --- | --- |
| version | R version 4.0.2 (2020-06-22) |
| os | CentOS Linux 7 (Core) |
| system | x86\_64, linux-gnu |
| ui | X11 |
| language | (EN) |
| collate | en\_US.UTF-8 |
| ctype | en\_US.UTF-8 |
| tz | America/New\_York |
| date | 2022-12-02 |

##### Code

```
options(yaml.eval.expr = TRUE)
set.seed(seed = 12345)
knitr::opts_chunk$set(echo = FALSE,warning = FALSE, message = FALSE)

library(tidyverse)
library(knitr)
library(ggplot2)
library(DESeq2)
library(circlize)
library(ComplexHeatmap)
library(openxlsx)
library(gt)

if(params$counts_format == "salmon") {
  library(tximport)
  }


# Set some mutable globals -----
design_formula <- as.formula(params$design_formula)
results_dir <- gsub(pattern = "/$",replacement = "",x = params$results_dir)
results_dir <- file.path(results_dir,params$counts_format)
check_dir(results_dir,is_dir = TRUE)
design_df <- readr::read_csv(file = params$design_file,col_names = TRUE)
contrast_col <- params$contrast_col

# Load annotation_df if specified -----


if(!is.null(params$annotation_file)){
  annotation_df <- readr::read_delim(file = params$annotation_file,delim = "\t",col_names = TRUE)
  
} else if(!is.null(params$gtf_file)){
  annotation_df <-parse_gtf_to_annotation_df(gtf_file = params$gtf_file)
  annotation_file <- gsub(x = basename(params$gtf_file),pattern = ".gtf.*$",replacement = "_tx2gene.tsv")
  annotation_file <- file.path(results_dir,annotation_file)
  message("Writing tx2gene transcript map that was generated to: ",annotation_file)
  readr::write_delim(annotation_df,annotation_file,delim = "\t",col_names = TRUE)
} else {
  stop("Either a gtf file or a transcript map must be supplied for this to work!")
}

if(!params$annotation_merge_col %in% colnames(annotation_df)){
    stop("Annotation file loaded, but the annotation_merge_col: '",params$annotation_merge_col,"' is not found in the column names! Columns are: ",colnames(annotation_df))
}

collapsed_annotation_df <- annotation_df %>%
    dplyr::select(gene_id,gene_name,gene_type) %>%
    distinct(gene_id,.keep_all = TRUE)


# Create the contrasts_df that is used for consistent parsing -----
if(length(params$contrasts) == 0){
  unique_contrast_categories <- unique(design_df[[params$contrast_col]])
  contrasts_df <-cross_df(list(
    "contrast" = params$contrast_col,
    "target" = unique_contrast_categories,
    "control" = unique_contrast_categories)) %>%
    dplyr::filter(target != control)
  } else if(length(params$contrasts) >= 0){
  contrasts_df <- map_df(params$contrasts,function(x){
    if(length(x) != 3) stop("All contrasts need to be a length of 3! They should be equivalent to c(contrast_column,target,control)")
    tibble(
      contrast = x[1],
      target = x[2],
      control = x[3])
  })
}

contrasts_df <- contrasts_df %>%
  mutate(
    contrast_label = glue::glue("{contrast}_{target}-vs-{control}"),
    contrast_dir = file.path(results_dir,contrast_label)
    )


if(params$counts_format == "salmon") {
    dds_dataset <- import_salmon(
      design_df = design_df,
      design_formula = design_formula,
      sf_file_col = params$sf_file_col,
      sample_id_col = params$sample_id_col,
      tx2gene = annotation_df,
      tximport_params = params$tximport_params,
      DESeqDataSetFromTximport_params = params$DESeqDataSetFromTximport_params
      )
    } else if(params$counts_format == "featureCounts") {
    dds_dataset <- import_featureCounts(
      design_df = design_df,
      featureCounts_file = params$counts_file,
      design_formula = design_formula,
      sample_id_col = params$sample_id_col,
      rowname_col = params$counts_rowname
    )
  }


dds <- do.call(
  DESeq2::DESeq,args = c(
    list("object" = dds_dataset),
    params$DESeq_params
    )
  )

filtered_colData <- SummarizedExperiment::colData(dds) %>%
  as.data.frame() %>%
  dplyr::select(!any_of(params$colData_scrub_filters))

# Collect vst counts from the DESeq2 object -----
vst_assay <- DESeq2::vst(dds,blind = FALSE)
vst_counts  <- SummarizedExperiment::assay(vst_assay)

message("Saving base dds object: ",file.path(results_dir,"DESeq2_object.RDS"))
saveRDS(object = dds,file = file.path(results_dir,"DESeq2_object.RDS"))

message("Saving VST counts: ",file.path(results_dir,"vst_counts.csv"))
readr::write_csv(vst_counts %>%
                   as_tibble(rownames = params$counts_rowname),
                 file.path(results_dir,"vst_counts.csv"),col_names = TRUE)
message("Saving normalized counts: ",file.path(results_dir,"normalized_counts.csv"))
readr::write_csv(x = DESeq2::counts(dds,normalized = TRUE) %>%
            as_tibble(rownames = params$counts_rowname),
          file.path(results_dir,"normalized_counts.csv"),
          col_names = TRUE)

design_df %>%
  gt(rowname_col = params$sample_id_col) %>%
  tab_header(title = "Sample Information") %>%
  opt_row_striping(row_striping = TRUE) %>%
  tab_options(container.width = px(800),table.layout = "auto")


if(params$counts_format == "salmon"){
  analysis_summary <- tribble(
  ~`Processing Step`,~Parameter,~Value,
  "DESeqDataSet Preparation","Design File",params$design_file,
  "DESeqDataSet Preparation","Counts Format",params$counts_format,
  "DESeqDataSet Preparation","Column with counts files",params$sf_file_col,
  "Comparison Conditions","Design Formula",paste(as.character(design_formula),collapse = ""),
  "Comparison Conditions","Contrast Column",params$contrast_col
)
} else if (params$counts_format == "featureCounts") {
  analysis_summary <- tribble(
  ~`Processing Step`,~Parameter,~Value,
  "DESeqDataSet Preparation","Count File",params$counts_file,
  "DESeqDataSet Preparation","Design File",params$design_file,
  "DESeqDataSet Preparation","Counts Format",params$counts_format,
  "Comparison Conditions","Design Formula",paste(as.character(design_formula),collapse = ""),
  "Comparison Conditions","Contrast Column",params$contrast_col
)
}


analysis_summary %>%
  gt(groupname_col = "Processing Step",rowname_col = "Parameter") %>%
  tab_header(title = "Summary of DESeq2 Inputs") %>%
  tab_style(
    style = cell_text(weight = "bold"),
    locations = cells_row_groups()
    ) %>%
  opt_row_striping(row_striping = TRUE) %>%
  tab_options(row.striping.include_stub = TRUE,column_labels.hidden = TRUE)


# Create PCA -----
if(length(params$pca_groups) == 0){
  intgroups <- names(filtered_colData)
  intgroups <- intgroups[which(intgroups != "sizeFactor")]
} else {
  intgroups <- params$pca_groups
}


pca_plot <- DESeq2::plotPCA(vst_assay,intgroup = intgroups)+
  common_theme(aspect.ratio = 1)

print(pca_plot)

export_ggplots(
  plot = pca_plot,
  plot_label = "DESeq2_pca_plot",
  output_dir = results_dir,
  additional_args = params$pca_plot_export
  )

# END -----


DESeq2::plotDispEsts(dds)

dsp_output_file_png <- file.path(results_dir,"DESeq2_dispersion_plot.png")
message("Saving Dispersion Plot (png): ",dsp_output_file_png)
png(filename = dsp_output_file_png,width = params$dispersion_export$width,height = params$dispersion_export$height,units = params$dispersion_export$units,res = params$dispersion_export$res)
DESeq2::plotDispEsts(dds)
dev.off()

dsp_output_file_pdf <- file.path(results_dir,"DESeq2_dispersion_plot.pdf")
message("Saving Dispersion Plot (pdf): ",dsp_output_file_pdf)
pdf(file = dsp_output_file_pdf,width = params$dispersion_export$width,height = params$dispersion_export$height)
DESeq2::plotDispEsts(dds)
dev.off()


DESeq2::plotSparsity(dds,normalized = TRUE)

sparsity_output_file_png <- file.path(results_dir,"DESeq2_sparsity_plot.png")
message("Saving sparsity Plot (png): ",sparsity_output_file_png)
png(filename = sparsity_output_file_png,width = params$sparsity_export$width,height = params$sparsity_export$height,units = params$sparsity_export$units,res = params$sparsity_export$res)
DESeq2::plotSparsity(dds,normalized = TRUE)
dev.off()

sparsity_output_file_pdf <- file.path(results_dir,"DESeq2_sparsity_plot.pdf")
message("Saving Dispersion Plot (pdf): ",sparsity_output_file_pdf)
pdf(file = sparsity_output_file_pdf,width = params$sparsity_export$width,height = params$sparsity_export$height)
DESeq2::plotSparsity(dds,normalized = TRUE)
dev.off()


generate_deseq_results_section <- function(
  dds,
  vst_counts,
  filtered_colData,
  contrast,
  target,
  control,
  contrast_label = NULL,
  output_folder = NULL,
  res_params = list(),
  indent_level = "###",
  annotation_df = NULL,
  annotation_merge_col = params$annotation_merge_col,
  res_merge_col = params$counts_rowname,
  volcano_color_params = params$volcano_plot_colors,
  volcano_export_params = params$volcano_plot_export,
  ma_color_params = params$ma_plot_colors,
  ma_export_params = params$ma_plot_export,
  if_export_params = params$if_plot_export,
  run_volcano = TRUE,
  run_ma = TRUE,
  run_if = TRUE,
  run_heatmap = TRUE,
  run_filtered_heatmap = TRUE
  ){
  if(!is.null(output_folder)) check_dir(output_folder,is_dir = TRUE)
  if (is.null(contrast_label)) contrast_label <- glue::glue("{contrast}_{target}-vs-{control}")
  dds_res <- create_dds_res(dds_obj = dds,contrast = contrast,target = target,control = control,contrast_label = contrast_label,output_folder = output_folder,res_params = res_params)
  results_df <- annotate_results_df(
    dds_res = dds_res,
    annotation_df = annotation_df,
    annotation_merge_col = annotation_merge_col,
    res_merge_col = res_merge_col)
  results_metadata <- S4Vectors::metadata(dds_res)
  
  addtl_deseq_opts <- list("DESeq2::DESeq" = params$DESeq_params,"DESeq2::results" = params$DESeq_results_params)
  for (i in names(addtl_deseq_opts)){
    if (length(addtl_deseq_opts[[i]]) == 0) addtl_deseq_opts[[i]] <- NULL 
  }
  
  if (length(addtl_deseq_opts) > 0){
    deseq_opts_table <- purrr::map_df(names(addtl_deseq_opts),function(x){
      tibble(
        Command = x,
        Option = names(addtl_deseq_opts[[x]]),
        Value = as.character(unlist(addtl_deseq_opts[[x]]))
      )
    })
  } else {
    deseq_opts_table <- tibble(
      Command = c(),
      Option = c(),
      Value = c()
    )
  }
  doc_params <- map_df(names(params),function(id){
  
  value_info <- params[[id]]
  value_names <- names(value_info)
  if(length(value_names) > 0){
    value_info <- paste(value_names,value_info,sep = " = ")
  }
  tibble(
    parameter = id,
    value = paste(value_info,collapse = ", ")
  )
})
  
  # save results -----
  ## RDS -----
  res_rds = file.path(output_folder,paste0(contrast_label,"_results.rds"))
  message("Writing DESeq2 results object: ",res_rds)
  saveRDS(object = dds_res,res_rds)
  ## csv -----
  res_csv <- file.path(output_folder,paste0(contrast_label,"_results.csv"))
  message("Writing DESeq2 results csv: ",res_csv)
  readr::write_csv(results_df,res_csv,col_names = TRUE)
  ## XLSX file -----
  res_xlsx = file.path(output_folder,paste0(contrast_label,"_results.xlsx"))
  message("Writing DESeq2 results xlsx: ",res_xlsx)
  wb <- openxlsx::write.xlsx(
    split(results_df,results_df$classification),
    file = res_xlsx,
    asTable = TRUE,
    overwrite = TRUE
    )
  ### DEG definitions -----
  addWorksheet(wb = wb,sheetName = "Classification Definitions")
  writeData(wb,sheet = "Classification Definitions",x = "Rules for classifying genes in DESeq2 results",colNames = TRUE,keepNA = TRUE,startRow = 1)
  writeDataTable(wb,sheet = "Classification Definitions",x = create_deg_descrip_table2(params$DESeq_results_params),colNames = TRUE,keepNA = TRUE,startRow = 2)
  ### DESeq2 summary -----
  addWorksheet(wb = wb,sheetName = "DESeq2 Summary")
  writeData(wb,sheet = "DESeq2 Summary",x = "DESeq2's summary output",colNames = TRUE,keepNA = TRUE,startRow = 1)
  writeData(wb,sheet = "DESeq2 Summary",x = capture.output(summary(dds_res)),colNames = TRUE,keepNA = TRUE,startRow = 2)
  ### Additional DESeq2 Params ----
  addWorksheet(wb = wb,sheetName = "DESeq2 Params")
  writeData(wb,sheet = "DESeq2 Params",x = "Additional parameters supplied to DESeq2",colNames = TRUE,keepNA = TRUE,startRow = 1)
  writeDataTable(wb,sheet = "DESeq2 Params",x = deseq_opts_table,colNames = TRUE,keepNA = TRUE,startRow = 2)
  ### Document Params ----
  addWorksheet(wb = wb,sheetName = "Rmd Params")
  writeData(wb,sheet = "Rmd Params",x = "Parameters used in the DESeq2 processing document",colNames = TRUE,keepNA = TRUE,startRow = 1)
  writeDataTable(wb,sheet = "Rmd Params",x = doc_params,colNames = TRUE,keepNA = TRUE,startRow = 2)
  saveWorkbook(wb,file = res_xlsx, overwrite = TRUE)
  
  # Volcano Plot -----
  if(run_volcano){
    
    volcano_plot <- create_volcano_plot(
    results_df = results_df,
    results_metadata = results_metadata,
    contrast = contrast,
    target = target,
    control = control,
    color_params = volcano_color_params
    )
  export_ggplots(
    plot = volcano_plot,
    plot_label = paste0(contrast_label,"_volcano_plot"),
    output_dir = output_folder,
    additional_args = volcano_export_params
  )
  }
  
  # MA Plot -----
  if(run_ma){
    ma_plot <- create_ma_plot(
    results_df = results_df,
    results_metadata = results_metadata,
    contrast = contrast,
    target = target,
    control = control,
    color_params = ma_color_params
    )
  export_ggplots(
    plot = volcano_plot,
    plot_label = paste0(contrast_label,"_ma_plot"),
    output_dir = output_folder,
    additional_args = ma_export_params
  )
  }
  
  # Independent Filter Plot -----
  if(run_if){
    if_plot <- create_if_plot(results_metadata = results_metadata,contrast = contrast,target = target,control = control)
  export_ggplots(
    plot = if_plot,
    plot_label = paste0(contrast_label,"_independent-filtering_plot"),
    output_dir = output_folder,
    additional_args = if_export_params
  )
  }
  if(run_heatmap){
    full_heatmap <- create_deg_heatmap(
      results_df = results_df,
      vst_counts = vst_counts,
      filtered_colData = filtered_colData,
      color_min = params$DEG_heatmap_min_color,
      color_mid = params$DEG_heatmap_mid_color,
      color_max = params$DEG_heatmap_max_color,
      rownames_col = params$counts_rowname)
    if(!is(full_heatmap,"character")){
      full_heat_file <- file.path(output_folder,paste0(contrast_label,"_full_deg_heatmap_plot"))
      message("Saving Full DEG Heatmap (png): ",paste0(full_heat_file,".png"))
      png(filename = paste0(full_heat_file,".png"),
          width = params$heatmap_export$width,
          height = params$heatmap_export$height,
          units = params$heatmap_export$units,
          res = params$heatmap_export$res)
      draw(full_heatmap)
      dev.off()
      message("Saving Full DEG Heatmap (pdf): ",paste0(full_heat_file,".pdf"))
      pdf(file = paste0(full_heat_file,".pdf"),
          width = params$heatmap_export$width,
          height = params$heatmap_export$height)
      draw(full_heatmap)
      dev.off()
    }
  }
  if(run_filtered_heatmap){
    filtered_indices <- which(filtered_colData[[contrast]] %in% c(target,control))
    filtered_samples <- row.names(filtered_colData)[filtered_indices]
    
    filtered_heatmap <- create_deg_heatmap(
      results_df = results_df,
      vst_counts = vst_counts,
      filtered_colData = filtered_colData,
      color_min = params$DEG_heatmap_min_color,
      color_mid = params$DEG_heatmap_mid_color,
      color_max = params$DEG_heatmap_max_color,
      rownames_col = params$counts_rowname,
      select_samples = filtered_samples)
    
    if(!is(filtered_heatmap,"character")){
      filtered_heat_file <- file.path(output_folder,paste0(contrast_label,"_filtered_deg_heatmap_plot"))
      message("Saving Filtered DEG Heatmap (png): ",paste0(filtered_heat_file,".png"))
      png(filename = paste0(filtered_heat_file,".png"),
          width = params$heatmap_export$width,
          height = params$heatmap_export$height,
          units = params$heatmap_export$units,
          res = params$heatmap_export$res)
      draw(filtered_heatmap)
      dev.off()
      message("Saving Filtered DEG Heatmap (pdf): ",paste0(filtered_heat_file,".pdf"))
      pdf(file = paste0(filtered_heat_file,".pdf"),
          width = params$heatmap_export$width,
          height = params$heatmap_export$height)
      draw(filtered_heatmap)
      dev.off()
    }
  }
  
  
  cat("  \n  \n")
  cat(glue::glue("  \n  \n{indent_level} {contrast}: {target} vs {control}  \n  \n"))
  print(summarize_deg_classification(results_df,contrast,target,control))
  cat("  \n  \n")
  print(summarize_top_degs(results_df,results_metadata,contrast,target,control))
  if(run_volcano) {
    cat("  \n  \n",glue::glue("{indent_level}# Volcano Plot  \n  \n"),sep = "\n");print(volcano_plot)
    }
  if(run_ma) {
    cat("  \n  \n",glue::glue("{indent_level}# MA Plot  \n  \n"),sep = "\n");print(ma_plot)
    }
  if(run_if) {
    cat("  \n  \n",glue::glue("{indent_level}# Independent Filtering Plot  \n  \n"),sep = "\n");print(if_plot)
    }
  if(run_heatmap) {
    cat("  \n  \n",glue::glue("{indent_level}# Full DEG Heatmap (all samples)  \n  \n"),sep = "\n");print(full_heatmap)
    }
  if(run_filtered_heatmap) {
    cat("  \n  \n",glue::glue("{indent_level}# Filtered DEG Heatmap (only contrast samples)  \n  \n"),sep = "\n");print(filtered_heatmap)
    }
  cat("  \n  \n")
  }

create_deg_descrip_table2(params$DESeq_results_params) %>%
  gt(rowname_col = "Classification") %>%
  tab_header(title = "Schema for classifying gene status") %>%
  opt_row_striping(row_striping = TRUE) %>%
  tab_options(row.striping.include_stub = TRUE,container.width = px(1000)) %>%
  tab_style(style = cell_text(style = "italic"),locations = cells_stub()) %>%
  cols_width(
    columns = c(padj,log2FoldChange)~px(200)
    )


purrr::pwalk(
    list(
      contrast = contrasts_df$contrast,
      target = contrasts_df$target,
      control = contrasts_df$control,
      contrast_label = contrasts_df$contrast_label,
      contrast_dir = contrasts_df$contrast_dir
      ),
    function(contrast,target,control,contrast_label,contrast_dir){
      generate_deseq_results_section(
      dds = dds,
      contrast = contrast,
      vst_counts = vst_counts,
      filtered_colData = filtered_colData,
      target = target,
      control = control,
      output_folder = contrast_dir,
      contrast_label = contrast_label,
      annotation_df = collapsed_annotation_df,
      annotation_merge_col = params$annotation_merge_col,
      res_merge_col = params$counts_rowname,
      indent_level = "###",
      res_params = params$DESeq_results_params,
      volcano_color_params = params$volcano_plot_colors,
      volcano_export_params = params$volcano_plot_export,
      ma_color_params = params$ma_plot_colors,
      ma_export_params = params$ma_plot_export,
      run_volcano = params$run_volcano,
      run_ma = params$run_ma,
      run_if = params$run_if,
      run_heatmap = params$run_heatmap,
      run_filtered_heatmap = params$run_filtered_heatmap
    )
    }
  )


summarize_deg_classification <- function(results_df,contrast,target,control,...){
  .args <- list(...)
  # Create the table to summarize how genes were classified
  results_df %>%
    group_by(classification) %>%
    summarize(
      Number = n()
    ) %>%
    ungroup() %>%
    mutate(Percentage = Number / sum(Number)) %>%
    gt(rowname_col = "classification",caption = "Summary of the classification for each gene after performing DESeq2 analysis.") %>%
    tab_header(title = "DESeq2 Results Summary",subtitle = glue::glue("{contrast}: {target} vs {control}")) %>%
    fmt_percent(columns = "Percentage",decimals = 2) %>%
    fmt_number(columns = "Number",decimals = 0) %>%
    summary_rows(columns = "Percentage",fns = list(Total = ~sum(.)),formatter = fmt_percent,decimals = 2) %>%
    summary_rows(columns = "Number",fns = list(Total = ~sum(.)),formatter = fmt_number,decimals = 0) %>%
    tab_style(style = cell_text(weight = "bold"),locations = list(cells_grand_summary(),cells_stub_grand_summary())) %>%
    opt_row_striping(row_striping = TRUE) %>%
    tab_options(row.striping.include_stub = TRUE) %>%
    cols_width(
      columns = c(Number,Percentage)~px(200)
    )
}

summarize_top_degs <- function(results_df,results_metadata,contrast,target,control,rowname_col = params$counts_rowname,table_width = px(1200),...){
  results_df %>%
    dplyr::filter(classification == "DEG") %>%
    dplyr::mutate(
      status = dplyr::case_when(
        log2FoldChange < 0 ~ "Downregulated",
        log2FoldChange == 0 ~ "No Change",
        log2FoldChange > 0 ~ "Upregulated"
      )
    ) %>%
    dplyr::group_by(status) %>%
    dplyr::arrange(padj,.by_group = TRUE) %>%
    dplyr::slice_head(n = 10) %>%
    dplyr::ungroup() %>%
    dplyr::arrange(padj,desc(log2FoldChange)) %>%
    gt(groupname_col = "status",rowname_col = params$counts_rowname,caption = "Top 10 up/down-regulated genes relative to control.") %>%
    tab_header(title = "Top Differentially Expressed Genes",subtitle = glue::glue("{contrast}: {target} vs {control}")) %>%
    fmt_scientific(columns = c("pvalue","padj"),decimals = 2) %>%
    fmt_number(columns = c("baseMean","log2FoldChange","lfcSE","stat"),decimals = 3) %>%
    tab_style(style = cell_text(weight = "bold"),locations = cells_row_groups()) %>%
    row_group_order(groups = c("Upregulated", "Downregulated")) %>%
    opt_row_striping(row_striping = TRUE) %>%
    tab_options(row.striping.include_stub = TRUE,table.layout = "auto",container.width = table_width) %>%
    tab_style(style = cell_text(size = "small"),locations = list(cells_body(),cells_stub(),cells_source_notes())) %>%
    cols_width(
      columns = c(pvalue,padj)~px(200),
      columns = c(log2FoldChange,baseMean,lfcSE,stat)~px(75)
    ) %>%
    data_color(
      columns = "log2FoldChange",alpha = 0.5,autocolor_text = FALSE,
      colors = scales::col_numeric(
        palette = c("#00FFFF","#FFFFFF","#E10022"),
        domain = c(min(results_df$log2FoldChange,na.rm = TRUE),max(results_df$log2FoldChange,na.rm = TRUE))
      )) %>%
    data_color(
      columns = "padj",alpha = 0.3,autocolor_text = FALSE,
      colors = scales::col_numeric(
        palette = c("#00ff00","#FFFFFF"),
        domain = c(0,results_metadata$alpha)
      )) %>%
    cols_hide("classification") %>%
    gt::tab_source_note(source_note = md("**baseMean**: The mean of the normalized counts between all samples")) %>%
    gt::tab_source_note(source_note = md("**log2FoldChange**: log~2~ fold change of the target / control")) %>%
    gt::tab_source_note(source_note = md("**lfcSE**: Standard error of the log~2~ fold change")) %>%
    gt::tab_source_note(source_note = md("**stat**: The statistic used to calculate the p-value")) %>%
    gt::tab_source_note(source_note = md("**pvalue**: p-value of the comparison, prior to adjustment")) %>%
    gt::tab_source_note(source_note = md("**padj**: The adjusted p-value, accounting for multiple comparisons"))
}

create_volcano_plot <- function(
  results_df,
  results_metadata,
  contrast,
  target,
  control,
  color_params = c()
  ){
  # Create Volcano Plots -----
  
  volcano_plot <- ggplot(results_df,aes(x = log2FoldChange,y = -log10(padj),color = classification))+
    ggplot2::geom_point()+
    scale_y_continuous(expand = expansion(mult = c(0,0.05)))+
    geom_hline(yintercept = -log10(results_metadata$alpha),linetype = 2)+
    geom_vline(xintercept = c(-results_metadata$lfcThreshold,results_metadata$lfcThreshold),linetype = 2)+
    ggtitle("Differentially Expressed Genes",glue::glue("{contrast}: {target} vs {control}"))+
    labs(
      x = expression(log[2](Fold~Change)),
      y = expression(-log[10](P[adj])),
    )+
    common_theme(aspect.ratio = 1)
  
  if(length(color_params) > 0) volcano_plot <- volcano_plot + scale_color_manual(values = color_params)
  
  volcano_plot
  
  
}

create_ma_plot <- function(
  results_df,
  results_metadata,
  contrast,
  target,
  control,
  color_params = c()
  ){
  # Create Volcano Plots -----
  
  ma_plot <- ggplot(results_df,aes(x = log2(baseMean),y = log2FoldChange,color = classification))+
    geom_point()+
    scale_y_continuous(expand = expansion(mult = c(0,0.05)))+
    labs(
      x = expression(log[2](Base~Mean)),
      y = expression(-log[2](Fold~Change)),
    )+
    ggtitle("Differentially Expressed Genes",glue::glue("{contrast}: {target} vs {control}"))+
    common_theme(aspect.ratio = 1)

  if(length(color_params) > 0) ma_plot <- ma_plot + scale_color_manual(values = color_params)
  
  ma_plot
  
  
}

create_if_plot <- function(results_metadata,contrast,target,control){
  annotation_text <- glue::glue("Filter Threshold: {round(results_metadata$filterThreshold,3)}\nFilter Theta: {round(results_metadata$filterTheta,3)}")
  
  if_plot <- ggplot(results_metadata$filterNumRej)+
    geom_point(aes(x = theta,y = numRej))+
    geom_line(data = as.data.frame(results_metadata$lo.fit),aes(x = x,y = y),size = 1)+
    geom_vline(xintercept = results_metadata$filterTheta)+
    labs(
      x = "Theta",
      y = "Number of Rejections",
    )+
    annotate(geom = "text",x = results_metadata$filterTheta+.02,y = 0.1,label = annotation_text,hjust = 0,vjust = 0)+
    ggtitle("DESeq2 Independent Filtering",glue::glue("{contrast}: {target} vs {control}"))+
    common_theme(aspect.ratio = 1)
  
  if_plot
  
}


create_deg_heatmap <- function(
  results_df,
  vst_counts,
  filtered_colData,
  color_min = params$DEG_heatmap_min_color,
  color_mid = params$DEG_heatmap_mid_color,
  color_max = params$DEG_heatmap_max_color,
  rownames_col = params$counts_rowname,
  select_samples = NULL
  ){
  sig_genes <- results_df %>%
    dplyr::filter(classification == "DEG") %>%
    dplyr::pull(rownames_col)
  
  if(length(sig_genes) > 0){
    message(length(sig_genes)," DEGs detected; we can run the heatmap!")
    sig_idx <- match(sig_genes,rownames(vst_counts))
    
    if(!is.null(select_samples)){
      vst_counts <- vst_counts[,select_samples]
      colData_indices <- which(row.names(filtered_colData) %in% select_samples)
      filtered_colData <- filtered_colData[colData_indices,]
      message("Filtering VST for specific samples: ",paste(select_samples,collapse = " "))
    }
    
    # Take the vst counts and filter for these indices
    # Then convert to z-score
    z_sig <- vst_counts[sig_idx,] %>%
      t() %>%
      scale() %>%
      t()
    
    # Collect anything in the design df
    th <- HeatmapAnnotation(df = filtered_colData,which = "column")
    # Plot with sensible defaults
    deg_heatmap <- Heatmap(
      z_sig,
      top_annotation = th,
      col = circlize::colorRamp2(
        breaks = c(min(z_sig),0,max(z_sig)),
        colors =c(params$DEG_heatmap_min_color,params$DEG_heatmap_mid_color,params$DEG_heatmap_max_color)
      ),
      cluster_rows = params$DEG_heatmap_cluster_rows,
      cluster_columns = params$DEG_heatmap_cluster_cols,
      border = "#000000",
      name = "z-score",
      show_row_names = params$DEG_heatmap_show_row_names,
      show_column_names = params$DEG_heatmap_show_col_names
    )
    return(deg_heatmap)
  } else{
    message("No DEGs detected; skipping heatmap!")
    return("No DEGs detected; skipping heatmap!")
  }
}


tribble(
  ~`File Type`,~`File Pattern`,~`Explanation`,
  "DESeq2 data object","DESeq2_object.RDS","An R-loadable object containing the object created by the DESeq() function.",
  "DESeq2 VST counts","vst_counts.csv","The variance-stabilized transformation counts output after using the DESeq() and vst() functions.",
  "DESeq2 normalized counts","normalized_counts.csv","The normalized counts output after using the DESeq() and counts(normalized = TRUE) functions.",
  "Dispersion plot","DESeq2_dispersion_plot.(png|pdf)","The dispersion plot output by DESeq2.",
  "Sparsity plot","DESeq2_sparsity_plot.(png|pdf)","The sparsity plot output by DESeq2.",
  "PCA plot","DESeq2_pca_plot.(png|pdf)","The PCA plot output by DESeq2.",
  "DEG Heatmap","<contrast folder>/<contrast>_deg_heatmap_plot.(png|pdf)","A heatmap of z-scored vst counts for the differentially expressed genes in the contrast.",
  "Independent Filtering plot","<contrast folder>/<contrast>_independent-filter_plot.(png|pdf)","The independent filtering plot for the contrast.",
  "MA plot","<contrast folder>/<contrast>_ma_plot.(png|pdf)","The MA plot for gene expression values in the contrast.",
  "Volcano plot","<contrast folder>/<contrast>_ma_plot.(png|pdf)","The volcano plot for gene expression values in the contrast.",
  "DESeq2 results object","<contrast folder>/<contrast>_results.rds","The results object calculated by DESeq2 for the contrast.",
  "DESeq2 results csv","<contrast folder>/<contrast>_results.csv","A csv file containing the results calculated by DESeq2 for the contrast. The classification column annotates gene status according to DESeq2's methodology.",
  "DESeq2 results xlsx","<contrast folder>/<contrast>_results.xlsx","An xlsx file containing the results calculated by DESeq2 for the contrast. Each tab contains the genes matching the classification according to DESeq2's methodology."
) %>%
  gt() %>%
  opt_row_striping(row_striping = TRUE) %>%
  tab_options(row.striping.include_stub = TRUE,table.layout = "auto",container.width = px(1000))


map_df(names(params),function(id){
  
  value_info <- params[[id]]
  value_names <- names(value_info)
  if(length(value_names) > 0){
    value_info <- paste(value_names,value_info,sep = " = ")
  }
  tibble(
    parameter = id,
    value = paste(value_info,collapse = ", ")
  )
}) %>%
  gt() %>%
  opt_row_striping(row_striping = TRUE) %>%
  tab_options(row.striping.include_stub = TRUE,table.layout = "auto",container.width = px(1000))


sessioninfo::package_info() %>%
  as_tibble() %>%
  dplyr::filter(attached == TRUE) %>%
  gt() %>%
  opt_row_striping(row_striping = TRUE) %>%
  tab_options(row.striping.include_stub = TRUE,table.layout = "auto",container.width = px(1000))

plat <- unlist(sessioninfo::platform_info())
tibble(
  Metric = names(plat),
  Value = plat
) %>%
  gt() %>%
  opt_row_striping(row_striping = TRUE) %>%
  tab_options(row.striping.include_stub = TRUE,table.layout = "auto",container.width = px(1000))

# Basic File Checks -----

#if(!file.exists(params$counts_file)) stop("counts_file not found!")
if(!file.exists(params$design_file)) stop("design_file not found!")

# Function arg checks -----
check_args(params$DESeq_params,func = DESeq2::DESeq,id = "DESeq2::DESeq")
check_args(params$DESeq_results_params,func = DESeq2::results,id = "DESeq2::results")


parse_gtf_to_annotation_df <- function(gtf_file){
  message("Parsing gtf file to a transcript map: ",gtf_file)
  read_delim(file = gtf_file,delim = "\t",comment = "#",col_names = F) %>%
    filter(X3 == "transcript") %>%
    mutate(
      transcript_id = str_extract(X9,pattern = '(?<=transcript_id ").*?(?=")'),
      gene_id = str_extract(X9,pattern = '(?<=gene_id ").*?(?=")'),
      gene_name = str_extract(X9,pattern = '(?<=gene_name ").*?(?=")'),
      gene_type = str_extract(X9,pattern = '(?<=gene_type ").*?(?=")')
    ) %>%
    dplyr::select(transcript_id,gene_id,gene_name,gene_type) %>%
    distinct(transcript_id,.keep_all = TRUE)
}

check_dir <- function(path,is_dir = FALSE,create = TRUE){
  if(!is_dir) path <- dirname(path)
  if(!dir.exists(path)){
    message(glue::glue("Directory doesn't exist: {path}"))
    if (create) {
      message(glue::glue("creating: {path}"))
      dir.create(path,recursive = TRUE)
    }
  }
}

check_args <- function(arg_list,func,id){
  if(length(arg_list) > 0){
    message("Validating additional parameters have been provided for: ",id)
    func_names <- names(formals(func))
    for(i in names(arg_list)){
      if(!i %in% func_names) stop(glue::glue("Found an unknown parameter. Please check: {i}"))
    }
    message("All parameters appear valid for: ",id)
  }
}


# ggplot2-related functions -----

common_theme <- function(...){
  ggplot2::theme(panel.background = element_rect(fill = NA),axis.line = element_line(colour = "#000000"),...)
}

export_ggplots <- function(plot,plot_label,output_dir,formats = c("png","pdf"),additional_args = list()){
  # Make sure dir exists
  check_dir(path = output_dir,is_dir = TRUE,create = TRUE)
  # Then save the appropriate format.
  for(i in formats){
    file_out <- file.path(output_dir,paste(plot_label,i,sep = "."))
    message(glue::glue("Saving {plot_label} ({i}): {file_out}"))
    do.call(ggsave,c(list(filename = file_out,plot = plot,device = i),additional_args))
  }
}

# END -----


import_salmon <- function(
  design_df,
  design_formula,
  sf_file_col,
  sample_id_col,
  tx2gene,
  tx2gene_cols = c("transcript_id","gene_id","gene_name"),
  tximport_params = list(),
  DESeqDataSetFromTximport_params = list()
  ){
  require(tximport)
  for(i in tx2gene_cols) {
    if(!i %in% colnames(tx2gene)) stop("Missing a required column in the annotation file for Salmon. The column is: ",i)
  }
  message("Processing counts from: Salmon")
  quant_files <- design_df[[sf_file_col]]
  names(quant_files) <- design_df[[sample_id_col]]
  
  # This part of the code is adapted from the vignette in the 'Tximport' package
  # https://bioconductor.org/packages/devel/bioc/vignettes/tximport/inst/doc/tximport.html#Salmon
  # Citation: Charlotte Soneson, Michael I. Love, Mark D. Robinson (2015): Differential analyses for RNA-seq: transcript-level estimates improve gene-level inferences. F1000Research
  
  tximport_params <- c(
    list(
      files = quant_files,
      type = "salmon",
      tx2gene = tx2gene
    ),
    tximport_params
  )
  
  txi_salmon <- do.call(tximport::tximport,tximport_params)
  
  deseq_design_df <- design_df %>%
    mutate(
      across(where(is.character),as.factor)
    )
  
  # Create DESeqDataSet from salmon txi -----
  DESeqDataSetFromTximport_params <- c(
    DESeqDataSetFromTximport_params,
    list(txi = txi_salmon,colData = deseq_design_df,design = design_formula)
  )
  dds_dataset <- do.call(DESeqDataSetFromTximport,DESeqDataSetFromTximport_params)
  
  dds_dataset
}


import_featureCounts <- function(
  design_df,
  featureCounts_file,
  design_formula,
  sample_id_col,
  rowname_col = params$counts_rowname
  ){

  message("Processing counts from: featureCounts")
  sample_order <- design_df[[sample_id_col]]
  if(length(sample_order) == 0) stop("The vector containing sample IDs is empty. Check that the 'sample_id_col' variable is correct.")
  
  counts_df <- readr::read_delim(file = featureCounts_file,delim = "\t",col_names = TRUE,comment = "#")
  if(!rowname_col %in% colnames(counts_df)) stop("counts_rowname parameter not found in featureCounts dataframe. Check this setting is correct.")
  counts_df_cols <- colnames(counts_df)
  
  updated_cols <- map_chr(counts_df_cols,function(x){
    sample_id <- str_extract_all(string = x,pattern = sample_order) %>% unlist()
    if(length(sample_id) == 0){
      return_sample_id <- x
    } else {
      sample_id <- unique(sample_id)
      id_lengths <- nchar(sample_id)
      return_sample_id <- sample_id[which(id_lengths == max(id_lengths))]
    }
    return_sample_id
  })
  
  colnames(counts_df) <- updated_cols
  purrr::walk2(counts_df_cols,updated_cols,function(.x,.y){
    message("remapped: ",.x," -> ",.y)
  })
  
  deseq_design_df <- design_df %>%
    mutate(
      across(where(is.character),as.factor)
    )
  
  
  ## Convert the appropriate column to rownames, remove unneeded columns, select all numeric cols, beautify column names, convert to matrix -----
  featureCounts_matrix <- counts_df %>%
    tibble::column_to_rownames(var = params$counts_rowname) %>%
    dplyr::select(-c(Chr,Start,End,Strand,Length)) %>%
    dplyr::select(where(is.numeric)) %>%
    dplyr::rename_with(basename) %>%
    dplyr::rename_with(~gsub(x = .x,pattern = "\\.bam$",replacement = "")) %>%
    as.matrix()

  featureCounts_matrix <- featureCounts_matrix[,sample_order]

  dds_dataset <- DESeqDataSetFromMatrix(countData = featureCounts_matrix,colData = deseq_design_df,design = design_formula)
  
  dds_dataset
}

annotate_results_df <- function(dds_res,annotation_df = NULL,annotation_merge_col = NULL,res_merge_col = NULL){
  results_metadata <- S4Vectors::metadata(x = dds_res)
  results_df <- tibble::as_tibble(x = dds_res,rownames = params$counts_rowname) %>%
    dplyr::mutate(
      classification = dplyr::case_when(
        baseMean == 0 & is.na(log2FoldChange) & is.na(pvalue) & is.na(padj) ~ "Zero Count",
        baseMean != 0 & !is.na(log2FoldChange) & is.na(pvalue) & is.na(padj) ~ "Outlier",
        baseMean != 0 & !is.na(log2FoldChange) & !is.na(pvalue) & is.na(padj) ~ "Low Count",
        padj < results_metadata$alpha ~ "DEG",
        TRUE ~ "Insignificant"
    )) %>%
    dplyr::arrange(padj)
  

  
  if(!is.null(annotation_df) && !is.null(annotation_merge_col) && !is.null(res_merge_col)){
    anno_cols <- colnames(annotation_df)
    anno_cols[which(anno_cols == annotation_merge_col)] <- res_merge_col
    colnames(annotation_df) <- anno_cols
    
    message("annotating results")
    results_df <- results_df %>%
      left_join(y = annotation_df,by = res_merge_col) %>%
      dplyr::select({{anno_cols}},everything())
  }
  
    results_df
}

create_deg_descrip_table2 <- function(results_params,alphaThreshold = 0.1){
  if(!"lfcThreshold" %in% names(results_params)) { lfcThreshold <- 0 }
  else { lfcThreshold <- results_params$lfcThreshold }
  
  lfcThreshold <- round(lfcThreshold,digits = 3)
  
  possible_alt_hypotheses <- c(
    "greaterAbs" = "|log2FC| > {lfcThreshold}",
    "lessAbs" = "|log2FC| < {lfcThreshold}",
    "greater" = "log2FC > {lfcThreshold}",
    "less" = "log2FC < -{lfcThreshold}"
  )
  
  if(!"altHypothesis" %in% names(results_params)) { altHypothesis <- "log2FC != {lfcThreshold}" }
  else if (!results_params$altHypothesis %in% names(possible_alt_hypotheses)) {stop("The specified altHypothesis is not recognized!")}
  else { altHypothesis <-  possible_alt_hypotheses[results_params$altHypothesis] }
  
  if("alpha" %in% names(results_params)) { alphaThreshold <- results_params$alpha }
  
  # Create a definitions table to add to the xlsx -----
  tribble(
    ~Classification,~`baseMean`,~`log2FoldChange`,~`pvalue`,~`padj`,~`Notes`,
    "Zero Count","= 0","NA","NA","NA","Zero count genes are removed from consideration prior to statistical analysis.",
    "Outlier","> 0","Exists","NA","NA","The gene is considered to be an outlier and is censured from analysis.",
    "Low Count","> 0","Exists","Exists","NA","An independent filtering step is performed prior to analysis that tries to establish a threshold for 'low count' genes. DESeq2's IF step tries to maximize the number of DEGs, so this may vary between contrasts.",
    "Insignificant","> 0","Exists","Exists",glue::glue(">= {alphaThreshold}"),"These are part of the actual analysis",
    "DEG (Differentially Expressed Gene)","> 0",glue::glue(altHypothesis),"Exists",glue::glue("< {alphaThreshold}"),"These are the genes that are considered to be of interest.",
  )
}


create_dds_res <- function(dds_obj,contrast,target,control,contrast_label = NULL,res_params = list(),output_folder = NULL,output_suffix = "_results.rds"){
  if (is.null(contrast_label)) contrast_label <- glue::glue("{contrast}_{target}-vs-{control}")
  message("Generating DESeq2 results for: ", contrast_label)
  dds_res_args <- c(
    list(
      "object" = dds_obj,
      "contrast" = c(contrast,target,control)
    ),
    res_params
    )
  dds_res <- do.call(DESeq2::results,args = dds_res_args)
  
  if(!is.null(output_folder)){
    check_dir(output_folder)
    res_rds = file.path(output_folder,paste0(contrast_label,output_suffix))
  message("Writing DESeq2 results object: ",res_rds)
  saveRDS(object = dds_res,res_rds)
  }
  dds_res

}
```
