## Supplementary material for "Human cytomegalovirus infection coopts chromatin organization to diminish TEAD1 transcription factor activity": html files for QC: QC_Report_UI_HFF_ATAC-seq.html

---

|  | general |
| --- | --- |
| Report generated at | 2022-02-06 21:59:06 |
| Title | CMV HS68 ATAC-seq UI |
| Description | CMV HS68 ATAC-seq UI |
| Pipeline version | v2.0.0 |
| Pipeline type | atac |
| Genome | hg19 |
| Aligner | bowtie2 |
| Sequencing endedness | {'rep1': {'paired\_end': True}, 'rep2': {'paired\_end': True}} |
| Peak caller | macs2 |

  

### Alignment quality metrics

---

#### SAMstat (raw unfiltered BAM)

|  | rep1 | rep2 |
| --- | --- | --- |
| Total Reads | 194571684 | 186494858 |
| Total Reads (QC-failed) | 0 | 0 |
| Duplicate Reads | 0 | 0 |
| Duplicate Reads (QC-failed) | 0 | 0 |
| Mapped Reads | 189121865 | 181106439 |
| Mapped Reads (QC-failed) | 0 | 0 |
| % Mapped Reads | 97.2 | 97.1 |
| Paired Reads | 194571684 | 186494858 |
| Paired Reads (QC-failed) | 0 | 0 |
| Read1 | 97285842 | 93247429 |
| Read1 (QC-failed) | 0 | 0 |
| Read2 | 97285842 | 93247429 |
| Read2 (QC-failed) | 0 | 0 |
| Properly Paired Reads | 186460626 | 178174908 |
| Properly Paired Reads (QC-failed) | 0 | 0 |
| % Properly Paired Reads | 95.8 | 95.5 |
| With itself | 187655484 | 179344202 |
| With itself (QC-failed) | 0 | 0 |
| Singletons | 1466381 | 1762237 |
| Singletons (QC-failed) | 0 | 0 |
| % Singleton | 0.8 | 0.8999999999999999 |
| Diff. Chroms | 121209 | 108234 |
| Diff. Chroms (QC-failed) | 0 | 0 |

  

#### Marking duplicates (filtered BAM)

|  | rep1 | rep2 |
| --- | --- | --- |
| Unpaired Reads | 0 | 0 |
| Paired Reads | 82074429 | 83363727 |
| Unmapped Reads | 0 | 0 |
| Unpaired Duplicate Reads | 0 | 0 |
| Paired Duplicate Reads | 19865232 | 24527946 |
| Paired Optical Duplicate Reads | 1623032 | 1751601 |
| % Duplicate Reads | 24.2039 | 29.4228 |

  

Filtered out (samtools view -F 1804):

- read unmapped (0x4)
- mate unmapped (0x8, for paired-end)
- not primary alignment (0x100)
- read fails platform/vendor quality checks (0x200)
- read is PCR or optical duplicate (0x400)

  

#### Fraction of mitochondrial reads (unfiltered BAM)

|  | rep1 | rep2 |
| --- | --- | --- |
| Rn = Number of Non-mitochondrial Reads | 188089164 | 179726560 |
| Rm = Number of Mitochondrial Reads | 4443423 | 5941320 |
| Rm/(Rn+Rm) = Frac. of mitochondrial reads | 0.023078810030221015 | 0.03199971906826318 |

  

#### SAMstat (filtered/deduped BAM)

|  | rep1 | rep2 |
| --- | --- | --- |
| Total Reads | 123418308 | 116551168 |
| Total Reads (QC-failed) | 0 | 0 |
| Duplicate Reads | 0 | 0 |
| Duplicate Reads (QC-failed) | 0 | 0 |
| Mapped Reads | 123418308 | 116551168 |
| Mapped Reads (QC-failed) | 0 | 0 |
| % Mapped Reads | 100.0 | 100.0 |
| Paired Reads | 123418308 | 116551168 |
| Paired Reads (QC-failed) | 0 | 0 |
| Read1 | 61709154 | 58275584 |
| Read1 (QC-failed) | 0 | 0 |
| Read2 | 61709154 | 58275584 |
| Read2 (QC-failed) | 0 | 0 |
| Properly Paired Reads | 123418308 | 116551168 |
| Properly Paired Reads (QC-failed) | 0 | 0 |
| % Properly Paired Reads | 100.0 | 100.0 |
| With itself | 123418308 | 116551168 |
| With itself (QC-failed) | 0 | 0 |
| Singletons | 0 | 0 |
| Singletons (QC-failed) | 0 | 0 |
| % Singleton | 0.0 | 0.0 |
| Diff. Chroms | 0 | 0 |
| Diff. Chroms (QC-failed) | 0 | 0 |

  

Filtered and duplicates removed

  

#### Fragment length statistics (filtered/deduped BAM)

|  | rep1 | rep2 |
| --- | --- | --- |
| Fraction of reads in NFR | 0.7299332753118654 | 0.6798476221265604 |
| Fraction of reads in NFR (QC pass) | True | True |
| Fraction of reads in NFR (QC reason) | OK | OK |
| NFR / mono-nuc reads | 3.889280246982018 | 3.171136467979799 |
| NFR / mono-nuc reads (QC pass) | True | True |
| NFR / mono-nuc reads (QC reason) | OK | OK |
| Presence of NFR peak | True | True |
| Presence of Mono-Nuc peak | True | True |
| Presence of Di-Nuc peak | False | True |

  


rep1


rep2

Open chromatin assays show distinct fragment length enrichments, as the cut
sites are only in open chromatin and not in nucleosomes. As such, peaks
representing different n-nucleosomal (ex mono-nucleosomal, di-nucleosomal)
fragment lengths will arise. Good libraries will show these peaks in a
fragment length distribution and will show specific peak ratios.

  

- NFR: Nucleosome free region

  

#### Sequence quality metrics (filtered/deduped BAM)

rep1


rep2

Open chromatin assays are known to have significant GC bias. Please take this
into consideration as necessary.

  

#### Annotated genomic region enrichment

|  | rep1 | rep2 |
| --- | --- | --- |
| Fraction of Reads in universal DHS regions | 0.5476839546366168 | 0.7708029232276763 |
| Fraction of Reads in blacklist regions | 0.006626237332633016 | 0.005049867882919886 |
| Fraction of Reads in promoter regions | 0.20512959876260822 | 0.3089813222635401 |
| Fraction of Reads in enhancer regions | 0.4093369599589714 | 0.47928862454643095 |

  

Signal to noise can be assessed by considering whether reads are falling into
known open regions (such as DHS regions) or not. A high fraction of reads
should fall into the universal (across cell type) DHS set. A small fraction
should fall into the blacklist regions. A high set (though not all) should
fall into the promoter regions. A high set (though not all) should fall into
the enhancer regions. The promoter regions should not take up all reads, as
it is known that there is a bias for promoters in open chromatin assays.

  

### Library complexity quality metrics

---

#### Library complexity (filtered non-mito BAM)

|  | rep1 | rep2 |
| --- | --- | --- |
| Total Fragments | 80214528 | 80900331 |
| Distinct Fragments | 62827542 | 60432815 |
| Positions with Two Read | 9880340 | 10269212 |
| NRF = Distinct/Total | 0.783244 | 0.747003 |
| PBC1 = OneRead/Distinct | 0.793932 | 0.76554 |
| PBC2 = OneRead/TwoRead | 5.048487 | 4.505091 |

  

Mitochondrial reads are filtered out by default.
The non-redundant fraction (NRF) is the fraction of non-redundant mapped reads
in a dataset; it is the ratio between the number of positions in the genome
that uniquely mapped reads map to and the total number of uniquely mappable
reads. The NRF should be > 0.8. The PBC1 is the ratio of genomic locations
with EXACTLY one read pair over the genomic locations with AT LEAST one read
pair. PBC1 is the primary measure, and the PBC1 should be close to 1.
Provisionally 0-0.5 is severe bottlenecking, 0.5-0.8 is moderate bottlenecking,
0.8-0.9 is mild bottlenecking, and 0.9-1.0 is no bottlenecking. The PBC2 is
the ratio of genomic locations with EXACTLY one read pair over the genomic
locations with EXACTLY two read pairs. The PBC2 should be significantly
greater than 1.

  

NRF (non redundant fraction)   
PBC1 (PCR Bottleneck coefficient 1)   
PBC2 (PCR Bottleneck coefficient 2)   
PBC1 is the primary measure. Provisionally

- 0-0.5 is severe bottlenecking
- 0.5-0.8 is moderate bottlenecking
- 0.8-0.9 is mild bottlenecking
- 0.9-1.0 is no bottlenecking

  

### Replication quality metrics

---

#### IDR (Irreproducible Discovery Rate) plots

rep1\_vs\_rep2


rep1-pr1\_vs\_rep1-pr2


rep2-pr1\_vs\_rep2-pr2


pooled-pr1\_vs\_pooled-pr2

#### Reproducibility QC and peak detection statistics

|  | overlap | idr |
| --- | --- | --- |
| Nt | 249269 | 175494 |
| N1 | 220606 | 144187 |
| N2 | 239776 | 166713 |
| Np | 258237 | 181543 |
| N optimal | 258237 | 181543 |
| N conservative | 249269 | 175494 |
| Optimal Set | pooled-pr1\_vs\_pooled-pr2 | pooled-pr1\_vs\_pooled-pr2 |
| Conservative Set | rep1\_vs\_rep2 | rep1\_vs\_rep2 |
| Rescue Ratio | 1.0359771973249783 | 1.0344684148745826 |
| Self Consistency Ratio | 1.0868970018947808 | 1.156227676558913 |
| Reproducibility Test | pass | pass |

  

Reproducibility QC

- N1: Replicate 1 self-consistent peaks (comparing two pseudoreplicates generated by subsampling Rep1 reads)
- N2: Replicate 2 self-consistent peaks (comparing two pseudoreplicates generated by subsampling Rep2 reads)
- Ni: Replicate i self-consistent peaks (comparing two pseudoreplicates generated by subsampling RepX reads)
- Nt: True Replicate consistent peaks (comparing true replicates Rep1 vs Rep2)
- Np: Pooled-pseudoreplicate consistent peaks (comparing two pseudoreplicates generated by subsampling pooled reads from Rep1 and Rep2)
- Self-consistency Ratio: max(N1,N2) / min (N1,N2)
- Rescue Ratio: max(Np,Nt) / min (Np,Nt)
- Reproducibility Test: If Self-consistency Ratio >2 AND Rescue Ratio > 2, then 'Fail' else 'Pass'

  

#### Number of raw peaks

|  | rep1 | rep2 |
| --- | --- | --- |
| Number of peaks | 298230 | 279233 |

  
Top 300000 raw peaks from macs2 with p-val threshold 0.01

### Peak calling statistics

---

#### Peak region size

|  | rep1 | rep2 | idr\_opt | overlap\_opt |
| --- | --- | --- | --- | --- |
| Min size | 150.0 | 150.0 | 150.0 | 150.0 |
| 25 percentile | 224.0 | 256.0 | 416.0 | 304.0 |
| 50 percentile (median) | 362.0 | 435.0 | 639.0 | 502.0 |
| 75 percentile | 655.0 | 755.0 | 967.0 | 823.0 |
| Max size | 3051.0 | 3079.0 | 3112.0 | 3112.0 |
| Mean | 496.87667236696507 | 561.0249504893762 | 735.0558435191663 | 619.3603317882411 |

  


rep1


rep2


idr\_opt


overlap\_opt

### Enrichment / Signal-to-noise ratio

---

#### Strand cross-correlation measures (filtered BAM)

|  | rep1 | rep2 |
| --- | --- | --- |
| Number of Subsampled Reads | 12500000 | 12500000 |
| Estimated Fragment Length | 0 | 0 |
| Cross-correlation at Estimated Fragment Length | 0.332118257469926 | 0.455608917004311 |
| Phantom Peak | 70 | 70 |
| Cross-correlation at Phantom Peak | 0.2564114 | 0.3337117 |
| Argmin of Cross-correlation | 1500 | 1500 |
| Minimum of Cross-correlation | 0.1054416 | 0.08308062 |
| NSC (Normalized Strand Cross-correlation coeff.) | 3.149784 | 5.483938 |
| RSC (Relative Strand Cross-correlation coeff.) | 1.50147 | 1.486361 |

  
  

Performed on subsampled (25000000) reads.
Such FASTQ trimming is for cross-corrleation analysis only.

- Normalized strand cross-correlation coefficient (NSC) = col9 in outFile
- Relative strand cross-correlation coefficient (RSC) = col10 in outFile
- Estimated fragment length = col3 in outFile, take the top value

  


rep1


rep2

#### TSS enrichment (filtered/deduped BAM)

|  | rep1 | rep2 |
| --- | --- | --- |
| TSS enrichment | 30.84435457452778 | 37.017616664136014 |

  


rep1


rep2

Open chromatin assays should show enrichment in open chromatin sites, such as
TSS's. An average TSS enrichment in human (hg19) is above 6. A strong TSS enrichment is
above 10. For other references please see https://www.encodeproject.org/atac-seq/

  

#### Jensen-Shannon distance (filtered/deduped BAM)

|  | rep1 | rep2 |
| --- | --- | --- |
| AUC | 0.14265999705489907 | 0.06127220296429397 |
| Synthetic AUC | 0.4966625690526387 | 0.49665085295427663 |
| X-intercept | 0.14599472869581215 | 0.25066891170365513 |
| Synthetic X-intercept | 0.0 | 0.0 |
| Elbow Point | 0.832102162514548 | 0.9165770118344378 |
| Synthetic Elbow Point | 0.5004257712904905 | 0.5056637859542541 |
| Synthetic JS Distance | 0.5400795166250758 | 0.698390137688027 |

  


### Peak enrichment

---

#### Fraction of reads in peaks (FRiP)

##### FRiP for macs2 raw peaks

|  | rep1 | rep2 | rep1-pr1 | rep2-pr1 | rep1-pr2 | rep2-pr2 | pooled | pooled-pr1 | pooled-pr2 |
| --- | --- | --- | --- | --- | --- | --- | --- | --- | --- |
| Fraction of Reads in Peaks | 0.470761947246919 | 0.7213752160767707 | 0.4692497647917844 | 0.7130975298334205 | 0.4658846238598572 | 0.7131630976019048 | 0.5957906538079868 | 0.5911454505155481 | 0.5893161178549225 |

  

##### FRiP for overlap peaks

|  | rep1\_vs\_rep2 | rep1-pr1\_vs\_rep1-pr2 | rep2-pr1\_vs\_rep2-pr2 | pooled-pr1\_vs\_pooled-pr2 |
| --- | --- | --- | --- | --- |
| Fraction of Reads in Peaks | 0.5819560109386579 | 0.44822948796219114 | 0.7073269913519872 | 0.5846836703514742 |

  

##### FRiP for IDR peaks

|  | rep1\_vs\_rep2 | rep1-pr1\_vs\_rep1-pr2 | rep2-pr1\_vs\_rep2-pr2 | pooled-pr1\_vs\_pooled-pr2 |
| --- | --- | --- | --- | --- |
| Fraction of Reads in Peaks | 0.544579369752843 | 0.4079985361653151 | 0.6594054295534816 | 0.548974678762894 |

  

For macs2 raw peaks:  

- repX: Peak from true replicate X
- repX-prY: Peak from Yth pseudoreplicates from replicate X
- pooled: Peak from pooled true replicates (pool of rep1, rep2, ...)
- pooled-pr1: Peak from 1st pooled pseudo replicate (pool of rep1-pr1, rep2-pr1, ...)
- pooled-pr2: Peak from 2nd pooled pseudo replicate (pool of rep1-pr2, rep2-pr2, ...)

  
For overlap/IDR peaks:  

- repX\_vs\_repY: Comparing two peaks from true replicates X and Y
- repX-pr1\_vs\_repX-pr2: Comparing two peaks from both pseudoreplicates from replicate X
- pooled-pr1\_vs\_pooled-pr2: Comparing two peaks from 1st and 2nd pooled pseudo replicates
